# NKX2.2 is required for timely NEUROG3 extinction and segregation of competing lineage programs in human pancreatic endocrine progenitors

**DOI:** 10.1101/2025.09.26.677825

**Authors:** Christopher Schaaf, Fiona M. Docherty, Madison X. Rodriguez, Patrick Sean McGrath, Christopher J. Hill, Kristen L. Wells, Lori Sussel

## Abstract

Pancreatic endocrine progenitors commit to the NEUROG3+ lineage before resolving into mono-hormonal cell types. The homeodomain factor NKX2.2 is required for islet formation in mice and humans, but its role at the endocrine progenitor stage is undefined. We generated NKX2.2-null human embryonic stem cells and differentiated them towards endocrine lineages. Mutant progenitors induced *NEUROG3* and formed endocrine committed cells at near wild-type frequency. Single cell transcriptomics showed persistent *NEUROG3* in mutant cells running α, β and δ programs, co-activation of the mutually exclusive *ARX* and *PAX4* networks, and no terminal lineage factor induction. Reanalysis of published islet NKX2.2 ChIP-seq showed that chromatin state at bound elements, not occupancy alone, predicted the direction of transcriptional change, including at *NEUROG3*, *NKX6.1*, *ARX* and *PAX4*. End-stage cultures yielded few insulin+ cells, with excess somatostatin, ghrelin, polyhormonal and hormone-negative cells. NKX2.2 is therefore dispensable for endocrine specification but required for resolution of endocrine fates.

## INTRODUCTION

In the developing mouse and human pancreas, a series of cell lineage decisions specify the distinct pancreatic cell types^1,2^. A pancreatic progenitor pool is delineated by the expression of the transcriptional regulator Pancreatic Duodenal homeobox 1 (PDX1)^3,4^, followed by the generation of tip cells marked by Pancreas Transcription Factor 1A (PTF1A)^5^, which ultimately give rise to the exocrine acinar tissue, or the bipotent progenitor trunk cells marked by SRY-box transcription factor 9 (SOX9)^3,6^. The bipotent progenitor population then gives rise to either ductal cells or endocrine progenitor cells (EP) marked by Neurogenin3 (NEUROG3)^3,7,8^. The EP cells give rise to the four distinct mature hormone-producing cell types: α cells (glucagon), β cells (insulin), δ cells (somatostatin) and PP cells (pancreatic polypeptide). Two additional transient endocrine cell populations are present during development: gastrin producing G-cells and ghrelin producing ε cells. The tightly coordinated paracrine and endocrine actions of the islet α, β and δ cells maintain proper glucose homeostasis, and disruption of this process presents clinically as diabetes.

The endocrine progenitor state is inherently transient. *NEUROG3* is expressed for a brief pulse in the endocrine progenitor population but becomes rapidly downregulated upon endocrine lineage specification^3,7^. Cells entering the endocrine lineage must then accomplish two tasks within a limited developmental window: establish the transcriptional program of a specific islet cell type and extinguish the progenitor program that enabled fate selection. These events are mechanistically distinct but often difficult to separate experimentally because loss-of-function studies typically evaluate endpoint cell populations. Consequently, loss of either a fate-specifying factor or a maturation factor can lead to the same apparent result: reduced numbers of the expected endocrine cell type. Evidence from mouse studies indicates that the competence of *NEUROG3*+ cells evolves as development proceeds^9^, implying that the length of time spent in the progenitor state itself carries instructive information for cell fate decisions.

In mice, several studies have characterized the transcription factor homeobox protein NK-2 homolog B (NKX2.2) as a necessary factor for α and β cell development^10–13^. NKX2.2 is expressed throughout the pancreas shortly after the onset of PDX1 expression and then becomes restricted to the α, β and PP lineages^14^. In mice deleted for *Nkx2.2*, β cells fail to form and there is a dramatic reduction in the number of α cells^10^, with a corresponding increase in the number of ghrelin-producing ε cells^11^. This defect is phenocopied in mice deleted for *Nkx2.2* specifically within the NEUROG3+ endocrine progenitor population^15^, which places its essential requirement downstream of endocrine specification. Molecular analyses have demonstrated that NKX2.2 binds and directly regulates target genes necessary for α and β cell fates^16,17^, and that it functions in cell-type specific manners within the islet, forming a repressor complex with GRG3 and DNMT3A to repress α cell factors in a β cell context^13^ and cooperating with the cofactor KLF4 to activate the α cell gene program in α cells^17^. Collectively, these studies establish that NKX2.2 is indispensable for endocrine cell fate specification and functions to coordinate broad transcriptional programs underlying endocrine differentiation and function.

In humans, homozygous mutations in *NKX2.2* led to undetectable serum insulin levels at birth and neonatal diabetes, suggesting regulatory functions similar to those observed in mice^18^. Glucagon expression in these patients was not reported, but increased serum ghrelin levels were reported in an independent case study^19^. Immunofluorescence analyses of NKX2.2 expression in human fetal pancreas tissue further suggested that, unlike in mice, *NKX2.2* expression is activated later during pancreas development, downstream of *NEUROG3*^3^. The human system therefore positions *NKX2.2* at the transition between endocrine cell specification and fate resolution, providing a unique opportunity to distinguish these two processes.

Directed differentiation of human pluripotent stem cells towards a β-like cell fate recapitulates key stages of pancreatic endocrine development in a tractable system that can be sampled throughout differentiation^20–26^. Beyond its potential application to cell replacement therapies for type 1 diabetes, this platform provides a powerful framework for investigating the early stages of human pancreas development at molecular resolution. However, a persistent limitation of current differentiation protocols is that, although endocrine cells are generated efficiently, they remain transcriptionally^22,27,28^ and functionally^29,30^ immature relative to primary human islets. Furthermore, differentiation products are heterogeneous, frequently containing polyhormonal endocrine cells and off-target populations^26,31–34^. The developmental step that is incompletely executed under these conditions remains unclear, as do the transcriptional mechanisms required to complete it. In this study, we used NKX2.2KO hESC lines to define the role of NKX2.2 in human endocrine development. We show that *NKX2.2* is induced concomitantly with *NEUROG3*, that endocrine lineage specification proceeds in its absence, and that loss of *NKX2.2* prevents cells from extinguishing the progenitor program and resolving a stable endocrine identity. Integration with human islet NKX2.2 occupancy data further reveals that the chromatin context predicts the transcriptional consequences of NKX2.2 binding and identifies the core fate-resolution circuitry under its direct control.

## RESULTS

### NKX2.2 is concomitantly expressed with NEUROG3 during the in vitro generation of stem cell derived β-like cells

To clarify when *NKX2.2* is expressed during the *in vitro* differentiation of hESC derived β-like cells, we differentiated the MEL-1 cell line expressing GFP from the *INS* promoter (MEL1-*INS*^GFP/w^, WT^35^) using the multi-step two-dimensional protocol developed by Hogrebe et al.^21^ (Figure 1A). Quantitative PCR (qPCR) was used to monitor transcription factor expression at each critical stage of the differentiation protocol and to identify when *NKX2.2* was activated. This analysis confirmed the differentiation process proceeded as previously described and indicated that *NKX2.2* expression was induced in the endocrine progenitor (EP) cells expressing *NEUROG3* from stage 5 day 1 (S5d1), reaching a peak in expression at S5d4/5 and maintained at a reduced level by the end of stage 5 (Figure 1B). Fluorescence microscopy analysis confirmed that these cells were successfully differentiated into β-like cells that robustly expressed GFP at the end stage of the 2D protocol (S6d14) and formed GFP-positive islet clusters upon 3D induction (Figure 1C).

**Figure 1.**
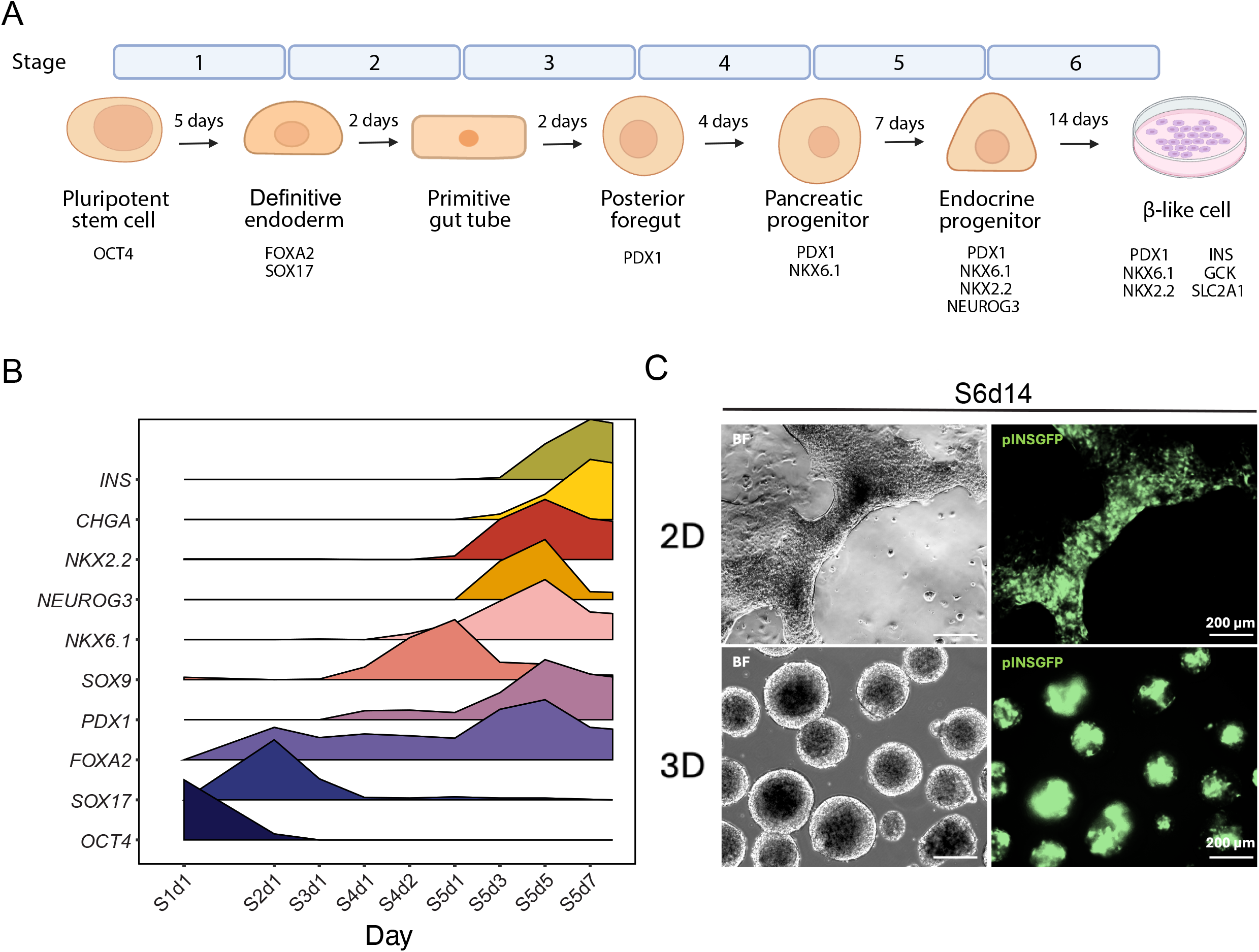
Directed differentiation of MEL-1 INS^GFP/w^ hESCs towards pancreatic endocrine cells. **(A)** Schematic of the six-stage differentiation protocol, adapted from Hogrebe et al.^21^ showing stage duration in days and the marker profile defining each intermediate: pluripotent stem cell (OCT4), definitive endoderm (FOXA2, SOX17), primitive gut tube, posterior foregut (PDX1), pancreatic progenitor (PDX1, NKX6.1), endocrine progenitor (PDX1, NKX6.1, NKX2.2, NEUROG3) and β-like cell (PDX1, NKX6.1, NKX2.2, INS, GCK, SLC2A1). Cells are cultured in two dimensions through stage 6 with optional aggregation into three-dimensional clusters at stage 6 day 7. **(B)** Quantitative PCR time course of stage-defining markers in WT cells across differentiation, from S1d1 to S5d7. Expression is shown as a density ridgeline per gene, with each gene scaled to its own maximum. *OCT4* is extinguished during stage 1 as *SOX17* and *FOXA2* rise. *PDX1* and *SOX9* rise through stages 3 and 4, and *NKX6-1* rises during stage 4 to peak at S5d1. *NEUROG3*, *NKX2-2*, *CHGA* and *INS* are induced during stage 5, with *NEUROG3* and *NKX2-2* rising together after S5d1 and peaking at S5d5. n = 3 independent WT differentiations. **(C)** Brightfield and pINSGFP images of WT cultures at S6d14 in two-dimensional culture (upper) and after aggregation into three-dimensional clusters (lower). GFP reports activity of the endogenous *INS* promoter. Scale bars, 200 μm.

This analysis showed that *NKX2.2* induction coincided with *NEUROG3* activation, aligning with the developmental timing inferred from human fetal tissue^3^ and distinct from the mouse, where *Nkx2.2* is expressed earlier in the pancreatic progenitor population^14^. Because *NKX2.2* is activated at the onset of the NEUROG3-defined progenitor state, it is likely to function in shaping or resolving that state rather than initiating it.

### Loss of NKX2.2 produces hormone-poor and polyhormonal cells at the end of the differentiation

To characterize the role of NKX2.2 in human endocrine cells, we utilized a CRISPR/Cas9^36^ targeting system to generate *NKX2.2* knockout (NKX2.2KO) hESC lines. A single guide RNA was used to target the first exon of *NKX2.2* in the WT parental cell line, which resulted in a homozygous single base-pair deletion at the gRNA cut site, a frameshift mutation, and an early stop codon 29 amino acids downstream of the cut site. The genetic mutations were confirmed by Sanger sequencing (Supplemental Figure 1A-C). To assess the ability of the mutant clones to differentiate through the multistep protocol, WT and NKX2.2KO cells from two independent clones (NKX2.2KOc3 and NKX2.2KOc5) were guided through the 2D differentiation protocol (Figure 1A). Bright field imaging demonstrated that both clones progressed through the entire protocol and produced confluent cultures at the mature β-like cell stage (S6d14) although there were some morphological differences observed between the two NKX2.2KO lines. Further, where the parent cell line showed robust GFP expression at S6d14, GFP was undetectable in both NKX2.2KO clones (Figure 2A). Immunofluorescent and confocal imaging at S6d14 confirmed a complete loss of the NKX2.2 protein in both clones and a substantial reduction in INS expression, with the residual INS signal restricted to sparse, scattered cells rather than the contiguous INS+ regions seen in WT (Figure 2B).

**Figure 2.**
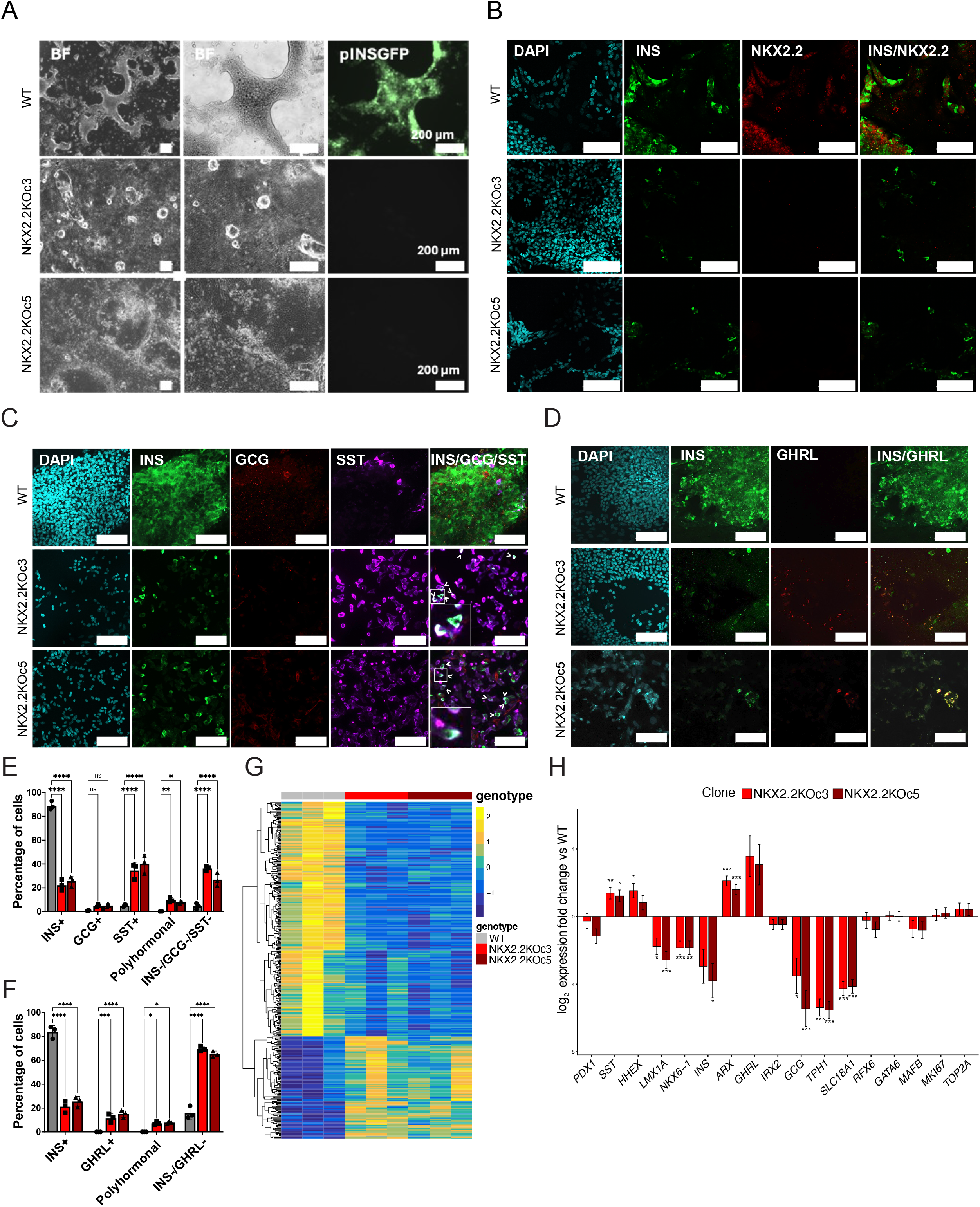
NKX2.2 is required for hormone expression at the end of the differentiation. All panels compare WT with the two independent NKX2.2 knockout clones, NKX2.2KOc3 and NKX2.2KOc5, at S6d14. **(A)** Brightfield and pINSGFP images at S6d14. Brightfield images are shown at two magnifications. WT cultures contain abundant GFP-positive cells, while neither knockout clone generates detectable pINSGFP signal. Scale bars, 200 μm. **(B)** Immunofluorescence for INS and NKX2.2 with DAPI at S6d14. NKX2.2 protein is undetectable in both knockout clones, confirming loss of the protein, and INS staining is sharply reduced. Scale bars, 100 μm. **(C)** Immunofluorescence for INS, GCG and SST with DAPI. Insets in the knockout rows show individual polyhormonal cells co-expressing INS and SST, and arrows mark polyhormonal cells. Scale bars, 100 μm. **(D)** Immunofluorescence for INS and GHRL with DAPI. Scale bars, 100 μm. **(E)** Quantification of the staining in (C), as the percentage of DAPI-positive cells that are INS+, GCG+, SST+, polyhormonal, or negative for all three hormones. INS+ cells fall from approximately 88% in WT to approximately 20 to 25% in the knockout clones, SST+ cells rise from approximately 4% to 35 to 40%, and hormone-negative cells rise from approximately 5% to 26 to 36%. Polyhormonal cells also increase in both clones. GCG+ cells are unchanged. **(F)** Quantification of the staining in (D), as the percentage of DAPI-positive cells that are INS+, GHRL+, polyhormonal, or negative for both. Cells expressing neither hormone make up 65 to 70% of the knockout cultures. For (E) and (F), bars are the mean of n = 3 independent differentiations with individual differentiations overlaid, scored across three slides per condition. Genotypes were compared by two-way ANOVA. *p < 0.05, **p < 0.01, ***p < 0.001, ****p < 0.0001, ns, not significant. **(G)** Heatmap of genes differentially expressed between WT and the NKX2.2 knockout clones at S6d14 (n = 396; padj < 0.05, |log2FC| > 1). Variance-stabilized counts were corrected for differentiation replicate by ComBat, filtered to genes with a mean normalized count of at least 5, and row z-scored. Rows are clustered; columns are ordered by genotype and are not clustered. Because the plotted genes were selected as differentially expressed against WT, the panel shows the expression pattern of that set and is not evidence about the relationship between the two clones. **(H)** Log2 fold change against WT for selected genes at S6d14, shown per clone. Bars are the fold change fitted by DESeq2 from a model that includes the differentiation replicate, and error bars are its standard error. Each clone was tested against WT separately. *p < 0.05, **p < 0.01, ***p < 0.001, Benjamini-Hochberg adjusted. n = 3 independent differentiations per clone.

Mice deleted for *Nkx2.2* display a complete absence of insulin-producing β cells, a severe reduction in glucagon-expressing α cells and a substantial increase in ghrelin-expressing ε cells, with somatostatin expression unaltered^10,11^. In humans, individuals with loss of function mutations in *NKX2.2* have no detectable C-peptide at birth and elevated serum ghrelin, while glucagon and somatostatin were not assessed^18,19^. To determine the hormone status of the cells differentiated from NKX2.2KO hESCs, we performed immunofluorescence for INS, glucagon (GCG) and somatostatin (SST), and separately for INS and ghrelin (GHRL), at S6d14 (Figure 2C-D). Only a small number of GCG-expressing cells were detectable in the WT parent cell line and in each of the NKX2.2KO clones, likely reflecting the bias of the differentiation protocol towards the β cell lineage. SST+ cells were substantially increased in both NKX2.2KO clones, and GHRL+ cells were increased, although not to the extent seen in mice lacking Nkx2.2. Polyhormonal cells, predominantly INS+/SST+ and INS+/GHRL+, were present in both NKX2.2KO lines.

Quantification confirmed and extended these observations (Figure 2E-F). Mono-hormonal INS+ cells fell from approximately 88% of WT cells to approximately 20% in both KO clones. GCG+ cells were unchanged between genotypes. SST+ cells rose from approximately 4% to approximately 35%, GHRL+ cells rose from approximately 1% to 11–15%, and polyhormonal cells rose in both scoring schemes. Cells expressing none of INS, GCG or SST rose from approximately 5% of WT cells to 25–38% of NKX2.2KO cells, and cells expressing neither INS nor GHRL made up the majority of the NKX2.2KO cultures at 65–72%. The differentiation therefore does not simply redistribute hormone output. A large fraction of NKX2.2KO cells reach the end of the protocol in a hormone-negative state.

To define the transcriptional basis of these changes, we performed bulk RNA sequencing and differential expression analysis at S6d14 of the WT and each NKX2.2KO cell line (n = 3 independent differentiations of WT, NKX2.2KOc3 and NKX2.2KOc5) and identified 396 genes that were significantly differentially expressed at this timepoint (Figure 2G). Consistent with the immunofluorescent analysis, *INS* and *NKX6-1* were strongly reduced, while *SST* and *ARX* were significantly increased. Although *GHRL* showed a large increase, it was not identified as significantly differentially expressed (Figure 2H). *HHEX* was also increased but not significantly. *PDX1* was unaffected. *IRX2* was unchanged and *GCG* was significantly reduced, suggesting that the elevation of *ARX* was not accompanied by induction of the α cell program. This likely reflects an increase in ARX+ progenitors, similar to what has been suggested by a recent single cell RNA-sequencing study on human embryonic pancreas tissue^37,38^. The enterochromaffin markers *LMX1A*, *TPH1* and *SLC18A1* were reduced, indicating that deletion of NKX2.2 suppresses the formation of this common off-target cell population^26,31^, which is consistent with its role in regulating the enterochromaffin population in the intestine^39,40^. *RFX6*, *GATA6*, *MAFB*, *MKI67* and *TOP2A* were largely unchanged at this stage. Overrepresentation analysis of the S6d14 differentially expressed genes returned neuroactive ligand-receptor interaction, insulin secretion, calcium signaling, and hormone signaling as the significantly enriched terms (Supplemental Figure 1D).

The end-stage phenotype is therefore not characterized by a loss of endocrine cells, but by the accumulation of endocrine cells that have altered hormone expression, are polyhormonal, or are hormone-negative. Although hormone production is altered, these cells do not converge on a distinct alternative fate. The remainder of this study investigates the developmental stage at which this cellular ambiguity originates.

### NKX2.2 is required for endocrine commitment

To identify the stage of differentiation at which NKX2.2 first becomes required, we performed flow cytometry for known stage-specific markers throughout the differentiation process in the WT and NKX2.2KOc5 cell lines. Flow cytometry for the pluripotency markers SOX2 and TRA-1-60 indicated that the NKX2.2KOc5 cell line maintained pluripotency after targeting (Figure 3A). At the posterior foregut stage (S4d1) there was no difference in the percentage of cells expressing PDX1 (Figure 3B). At the pancreatic progenitor stage (S5d1) the percentage of PDX1-expressing cells was marginally but significantly increased in the NKX2.2KO line, while the percentage of cells co-expressing PDX1 and NKX6.1 was unchanged (Figure 3C). By the end of the EP stage (S6d1), a clear separation emerged. The PDX1+ population remained unchanged in the NKX2.2KO cells, whereas the PDX1+/NKX6.1+ population was significantly decreased, indicating that in the absence of NKX2.2, *NKX6.1* expression is not maintained. The proportion of CHGA+ cells was significantly reduced, though the majority of cells in both lines remained CHGA+. Against that near-normal endocrine background, the number of INS+ cells was strongly reduced (Figure 3D).

**Figure 3.**
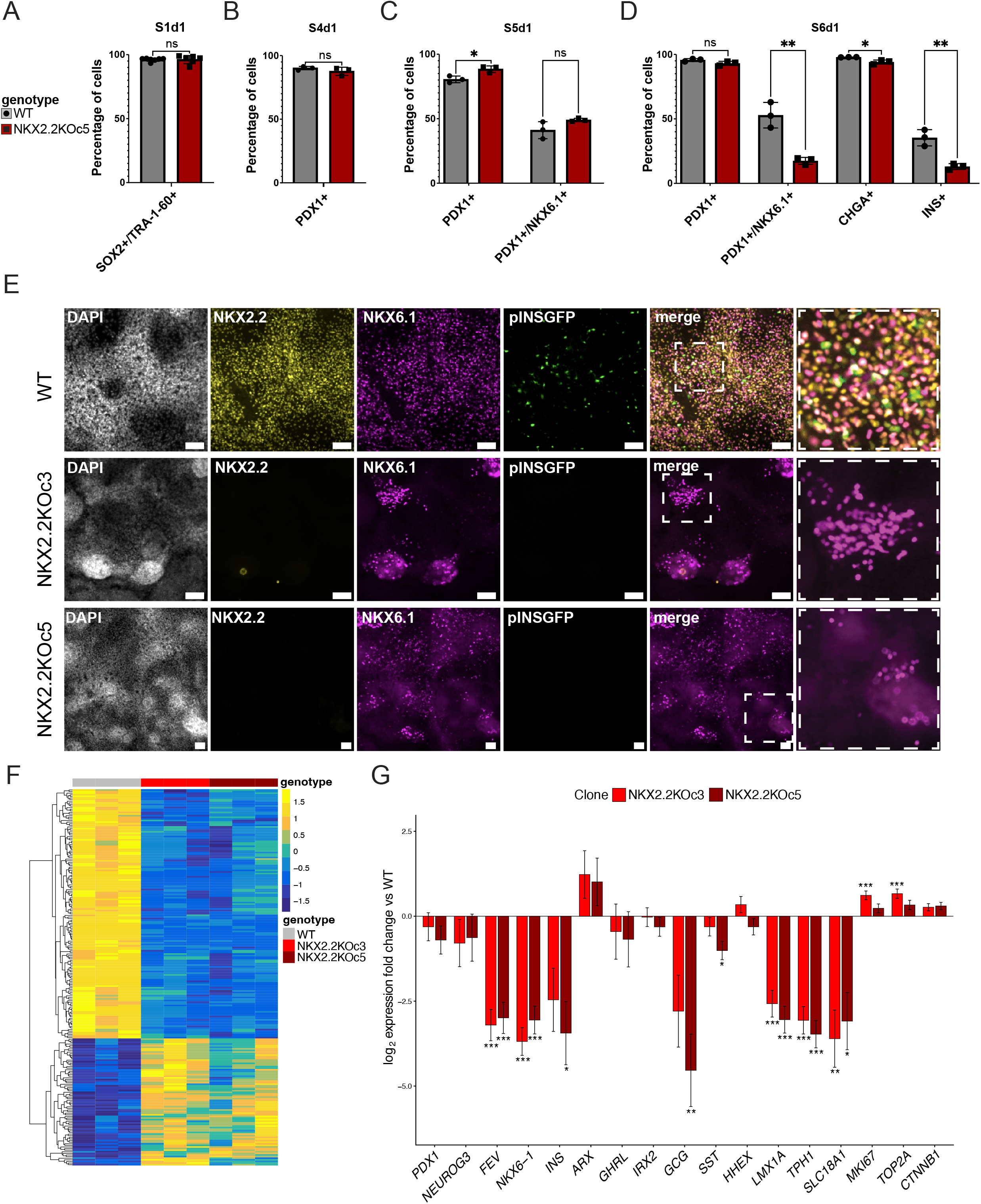
NKX2.2 is first required during the endocrine progenitor stage. (A to. **D)** Flow cytometry for stage-specific markers in WT and NKX2.2KOc5 cultures: (A) SOX2 and TRA-1-60 at S1d1, (B) PDX1 at S4d1, (C) PDX1 and PDX1/NKX6.1 at S5d1, and (D) PDX1, PDX1/NKX6.1, CHGA and INS at S6d1. Pluripotency and early pancreatic specification are unaffected. At S5d1 the PDX1+ fraction is slightly but significantly higher in the knockout. By S6d1 the PDX1+/NKX6.1+ and INS+ fractions are strongly reduced while CHGA+ cells are only slightly reduced, placing the first requirement for NKX2.2 within the endocrine progenitor stage. Bars are the mean of n = 3 independent differentiations with individual differentiations overlaid, compared by unpaired two-tailed t-test. *p < 0.05, **p < 0.01, ns, not significant. **(E)** Immunofluorescence for NKX2.2, NKX6.1 and pINSGFP with DAPI at S5d4, the middle of the endocrine progenitor stage. Boxed regions are shown at higher magnification at right. NKX2.2 protein is absent from both knockout clones, NKX6.1-positive cells are present but reduced and clustered, and pINSGFP is not detected. Scale bars, 200 μm. **(F)** Heatmap of genes differentially expressed between WT and the NKX2.2 knockout clones at S5d4 (n = 308; padj < 0.05, |log2FC| > 1), processed as in Figure 2G. **(G)** Log2 fold change against WT for selected genes at S5d4. Bars are the fold change fitted by DESeq2 from a model that includes the differentiation replicate, and error bars are its standard error. Each clone was tested against WT separately. *p < 0.05, **p < 0.01, ***p < 0.001, Benjamini-Hochberg adjusted. n = 3 independent differentiations per clone.

Immunofluorescence at S5d4, the middle of the EP stage, confirmed the phenotype suggested by the flow cytometry analysis (Figure 3E). NKX2.2 protein was absent from both KO clones, as expected. NKX6.1 protein was still detectable in both clones but was confined to discrete patches rather than distributed broadly across the culture as in WT, and pINSGFP was nearly absent. S5d4 therefore captures the phenotype at its onset, and we used this timepoint for all subsequent transcriptional analyses.

To define the transcriptional changes at this stage, we performed bulk RNA sequencing at S5d4 in the WT and NKX2.2KO cell lines (n = 3 independent differentiations each of WT, NKX2.2KOc3 and c5). Differential gene expression analysis revealed 308 significantly differentially expressed genes between the WT and both NKX2.2KO clones (Figure 3F). Consistent with the flow cytometry analysis, *PDX1* and *NEUROG3* expression were unaffected in the NKX2.2KO cell lines (Figure 3G). Despite preserved *PDX1* and *NEUROG3* expression, *FEV*, a marker of the intermediate EP population^41^, was strongly reduced, as were *NKX6-1* and *INS*. *ARX* was increased, although not significantly, *GHRL* was not significantly affected, *IRX2* was unchanged and *GCG* was significantly reduced. *SST* and *HHEX* were not elevated at S5d4, in contrast to their marked elevation at the terminal stage of differentiation (Figure 2H). The shift toward a δ cell-like hormone output of the NKX2.2KO cultures therefore develops after the onset of the primary defect and is not a feature of the initial disruption in endocrine differentiation. Markers of the enterochromaffin lineage, *LMX1A*, *TPH1* and *SLC18A1* were also reduced at this stage, indicating that loss of this lineage is already apparent at the progenitor stage rather than arising during endocrine maturation. *MKI67* and *TOP2A* were increased, with the increase in *TOP2A* reaching significance, consistent with persistence of a proliferative progenitor character rather than progression into a post-mitotic endocrine state; both returned to wild-type levels by S6d14 (Figure 2H). β-catenin (*CTNNB1*) was also modestly but significantly increased at S5d4 (Figure 3G), potentially suggesting a mechanism for failed endocrine progression.

The transcriptional pattern at S5d4 is particularly informative in that *NEUROG3* was induced normally, whereas multiple downstream targets failed to do so, reinforcing the conclusion that *NKX2.2* functions downstream of endocrine progenitor specification but before the establishment of differentiated endocrine identity. Overrepresentation analysis of the S5d4 differentially expressed genes returned neuroactive ligand-receptor interaction, insulin secretion, calcium signaling, serotonergic synapse and cAMP signaling as the significantly enriched terms (Supplemental Figure 1E). All eight genes driving the insulin secretion term were reduced in the NKX2.2KO. We interpret this as likely early signals of dysregulated β cell commitment due to these samples being assayed at the endocrine progenitor stage.

### Single cell transcriptomics localize the defect to endocrine lineage fate determination

Because β-like cell differentiation protocols generate heterogeneous populations^26^, we performed single cell RNA sequencing (scRNA-seq) on WT and NKX2.2KOc5 cells at S5d4 (n = 3 independent differentiations per genotype) corresponding to the mid-EP stage at which phenotypic differences were first observed. After quality filtering and doublet removal, 22,903 cells were retained. Clusters were annotated first on WT cells alone (Supplemental Figure 2A-B) and then by mapping of the WT annotations onto the NKX2.2KO cells. This resulted in 13 populations identified using previously described marker genes (Figure 4A, Table 1).

**Figure 4.**
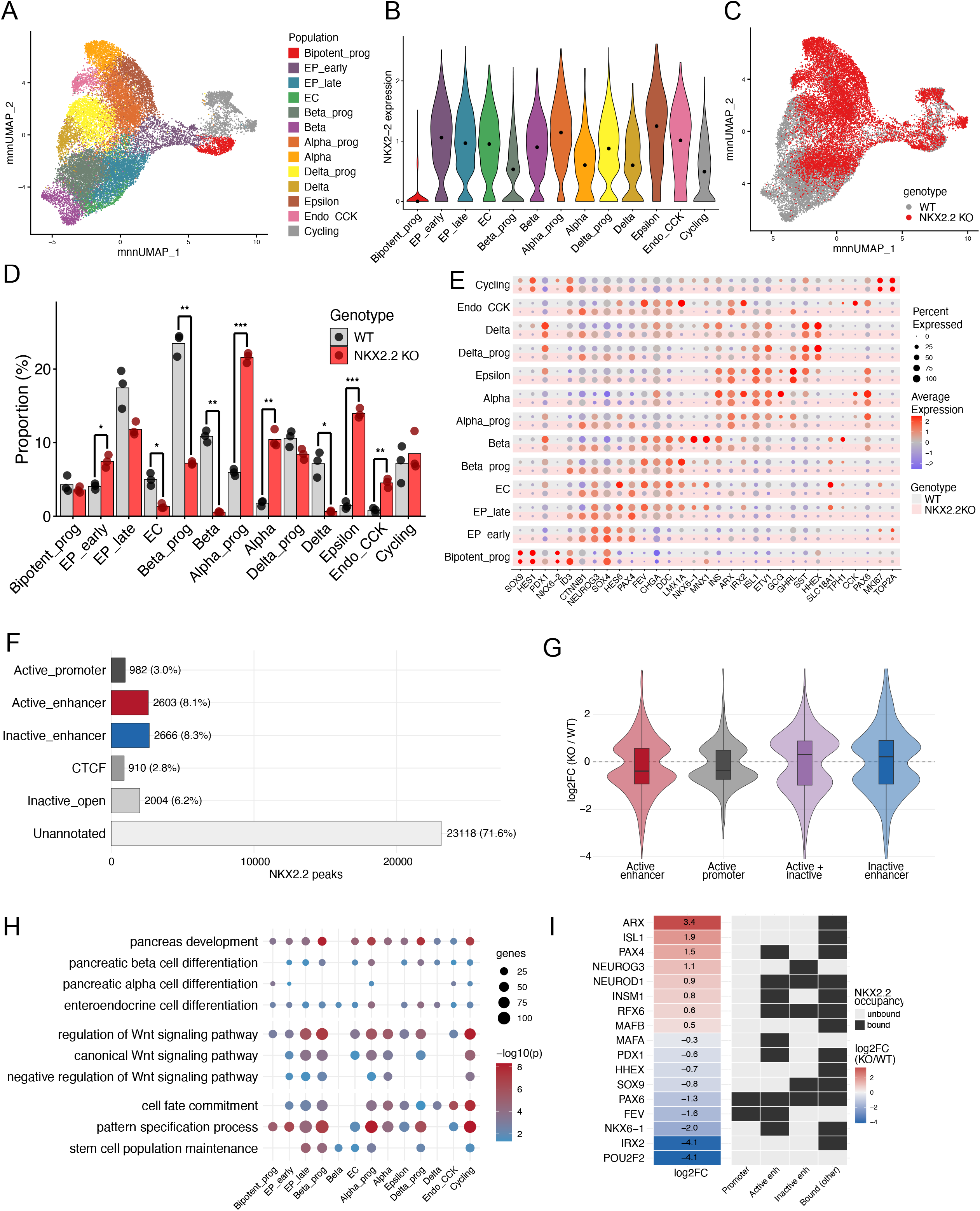
Single cell transcriptomics at S5d4 and NKX2.2 occupancy in human islets. (A to E, G to. **I)** Single cell RNA sequencing of WT and NKX2.2KOc5 cultures at S5d4, n = 3 independent differentiations per genotype, 22,903 cells after filtering. Cells of both genotypes were clustered together on the batch-corrected embedding, so cells of either genotype in the same cluster share a transcriptional neighborhood but are not assumed to be equivalent. **(F-I)** NKX2.2 ChIP-seq peaks are from adult human islets (Pasquali, Nat Genet, 2014) and were assigned to human islet regulome classes from the same study. **(A)** UMAP of all cells colored by annotated population. Thirteen populations were annotated from enriched expression of known markers (Table 1, Supplemental Figure 2). **(B)** Violin plots of *NKX2-2* expression across the annotated populations. Expression is low in bipotent progenitors and rises with endocrine commitment. *NKX2-2* transcript remains detectable in the knockout, since the edit is a single base-pair deletion rather than a deletion of the locus, so transcript level does not report protein. **(C)** The same UMAP colored by genotype. WT and NKX2.2KO cells occupy the same progenitor territory and diverge downstream of it. **(D)** Population proportions by genotype, as the percentage of each replicate’s cells assigned to each population. Points are individual differentiations. Genotypes were compared per population by Welch’s two-tailed t-test, Benjamini-Hochberg adjusted across all 13 populations. *p < 0.05, **p < 0.01, ***p < 0.001. **(E)** Dot plot of marker expression across populations, split by genotype. Dot size is the percentage of cells expressing the gene and color is mean scaled expression. **(F)** Distribution of the 32,283 reproducible NKX2.2 ChIP-seq peaks across human islet regulome classes. The majority fall outside annotated regulatory elements. **(G)** Distribution of pseudobulk log2 fold changes (NKX2.2KO over WT) for genes grouped by the chromatin state of their NKX2.2-bound elements. Active + inactive denotes genes with bound elements in both active and inactive classes. Genes bound at active enhancers (median log2FC −0.39, n = 581) or active promoters (−0.38, n = 259) shift down; genes bound at inactive enhancers (+0.21, n = 752) or at both active and inactive elements (+0.31, n = 368) shift up. Active versus inactive enhancer, two-sided Wilcoxon rank sum test, p = 0.0065. **(H)** Gene Ontology Biological Process enrichment for genes that are both NKX2.2-bound and differentially expressed, tested per population. Dot size is the number of genes in the term and color is −log10 of the adjusted p-value. Populations in which a term was not significantly enriched are blank rather than plotted as a non-significant point. Terms are grouped as pancreatic endocrine, WNT signaling and fate commitment. **(I)** Transcription factors defining the endocrine progenitor and islet lineages. Left, pseudobulk log2 fold change (NKX2.2KO over WT) across all cells. Right, NKX2.2 occupancy at each locus by chromatin class: promoter, active enhancer, inactive enhancer, or bound at an unannotated site. *NEUROG3* is bound only at an inactive enhancer and is upregulated; *NKX6-1* and *FEV* are bound at active elements and are downregulated; *ARX* and *IRX2* are bound only at unannotated sites; *POU2F2* is not bound.

**Table 1.**
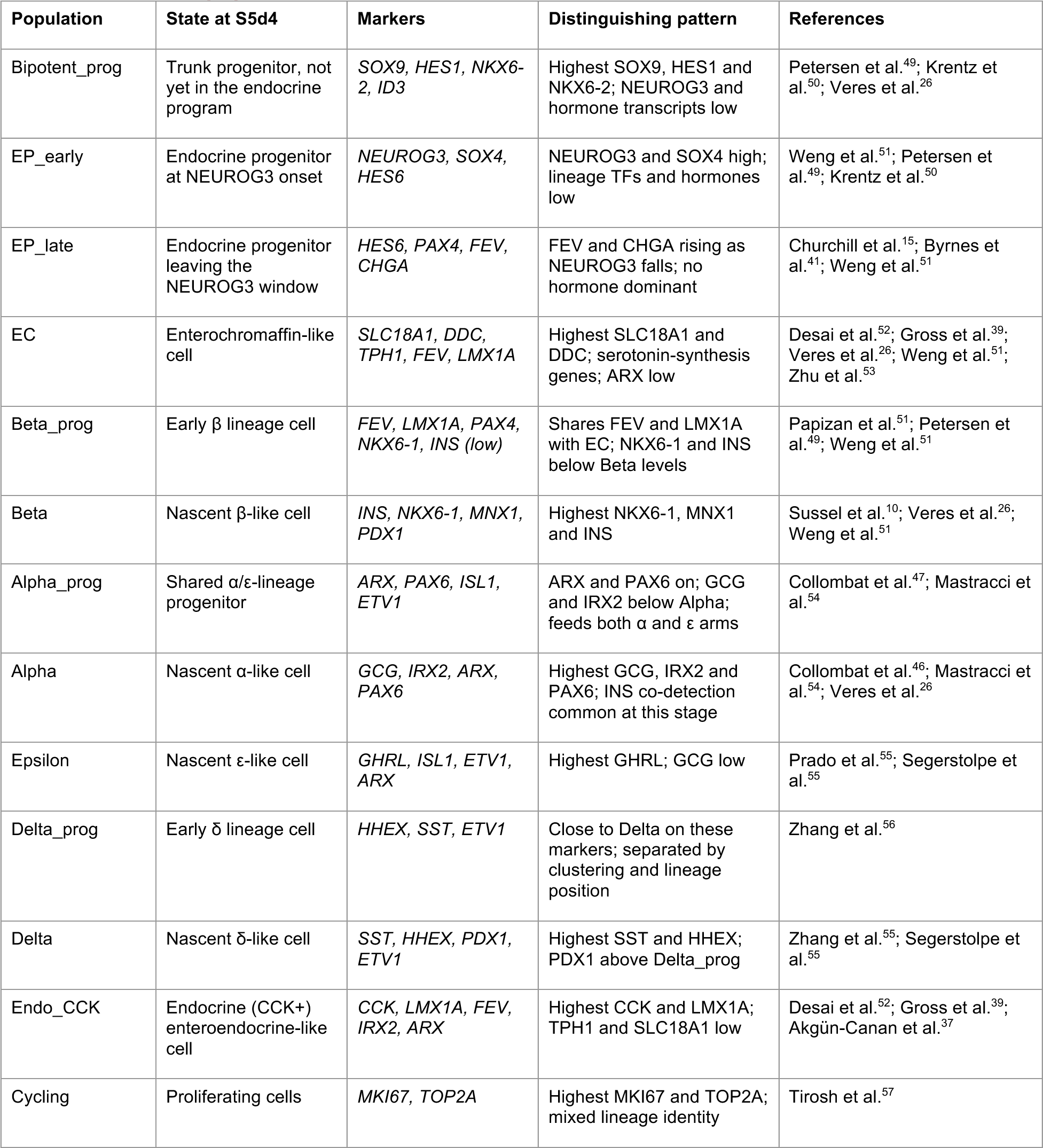
Annotated populations and canonical markers.

| Population | State at S5d4 | Markers | Distinguishing pattern | References |
|---|---|---|---|---|
| Bipotent_prog | Trunk progenitor, not yet in the endocrine program | *SOX9, HES1, NKX6-2, ID3* | Highest SOX9, HES1 and NKX6-2; NEUROG3 and hormone transcripts low | Petersen et al.<sup>49</sup>; Krentz et al.<sup>50</sup>; Veres et al.<sup>26</sup> |
| EP_early | Endocrine progenitor at NEUROG3 onset | *NEUROG3, SOX4, HES6* | NEUROG3 and SOX4 high; lineage TFs and hormones low | Weng et al.<sup>51</sup>; Petersen et al.<sup>49</sup>; Krentz et al.<sup>50</sup> |
| EP_late | Endocrine progenitor leaving the NEUROG3 window | *HES6, PAX4, FEV, CHGA* | FEV and CHGA rising as NEUROG3 falls; no hormone dominant | Churchill et al.<sup>15</sup>; Byrnes et al.<sup>41</sup>; Weng et al.<sup>51</sup> |
| EC | Enterochromaffin-like cell | *SLC18A1, DDC, TPH1, FEV, LMX1A* | Highest SLC18A1 and DDC; serotonin-synthesis genes; ARX low | Desai et al.<sup>52</sup>; Gross et al.<sup>39</sup>; Veres et al.<sup>26</sup>; Weng et al.<sup>51</sup>; Zhu et al.<sup>53</sup> |
| Beta_prog | Early β lineage cell | *FEV, LMX1A, PAX4, NKX6-1, INS (low)* | Shares FEV and LMX1A with EC; NKX6-1 and INS below Beta levels | Papizan et al.<sup>51</sup>; Petersen et al.<sup>49</sup>; Weng et al.<sup>51</sup> |
| Beta | Nascent β-like cell | *INS, NKX6-1, MNX1, PDX1* | Highest NKX6-1, MNX1 and INS | Sussel et al.<sup>10</sup>; Veres et al.<sup>26</sup>; Weng et al.<sup>51</sup> |
| Alpha_prog | Shared α/ε-lineage progenitor | *ARX, PAX6, ISL1, ETV1* | ARX and PAX6 on; GCG and IRX2 below Alpha; feeds both α and ε arms | Collombat et al.<sup>47</sup>; Mastracci et al.<sup>54</sup> |
| Alpha | Nascent α-like cell | *GCG, IRX2, ARX, PAX6* | Highest GCG, IRX2 and PAX6; INS co-detection common at this stage | Collombat et al.<sup>46</sup>; Mastracci et al.<sup>54</sup>; Veres et al.<sup>26</sup> |
| Epsilon | Nascent ε-like cell | *GHRL, ISL1, ETV1, ARX* | Highest GHRL; GCG low | Prado et al.<sup>55</sup>; Segerstolpe et al.<sup>55</sup> |
| Delta_prog | Early δ lineage cell | *HHEX, SST, ETV1* | Close to Delta on these markers; separated by clustering and lineage position | Zhang et al.<sup>56</sup> |
| Delta | Nascent δ-like cell | *SST, HHEX, PDX1, ETV1* | Highest SST and HHEX; PDX1 above Delta_prog | Zhang et al.<sup>55</sup>; Segerstolpe et al.<sup>55</sup> |
| Endo_CCK | Endocrine (CCK+) enteroendocrine-like cell | *CCK, LMX1A, FEV, IRX2, ARX* | Highest CCK and LMX1A; TPH1 and SLC18A1 low | Desai et al.<sup>52</sup>; Gross et al.<sup>39</sup>; Akgün-Canan et al.<sup>37</sup> |
| Cycling | Proliferating cells | *MKI67, TOP2A* | Highest MKI67 and TOP2A; mixed lineage identity | Tirosh et al.<sup>57</sup> |

In WT cells, *NKX2.2* expression was low in the bipotent progenitors, became upregulated in the EP_early population, and remained elevated throughout the committed endocrine populations (Figure 4B), matching the qPCR time course. WT and NKX2.2KO cells occupied the same progenitor territory but diverged into two distinct branches downstream of the shared progenitor clusters (Figure 4C).

Quantification of cell-type proportions across the single-cell dataset revealed that the genotype-dependent differences were highly concentrated within the committed progenitor and immature endocrine cell populations (Figure 4D). In the NKX2.2KO, there were significant reductions in enterochromaffin (EC) cells (EC; 1.42% in the KO versus 5.02% in the WT, adjusted p-val = 0.036), β progenitors (Beta_prog; 7.55% in the KO versus 23.65% in the WT, adjusted p-val <0.001) immature β-like cells (Beta; 0.57% in the KO versus 10.95% in the WT, adjusted p-val <0.001) and immature δ-like cells (Delta; 0.66% in the KO versus 7.20% in the WT, adjusted p-val = 0.036). In contrast, NKX2.2KO cells were significantly enriched in α progenitors (Alpha_prog; 22.60% in the KO versus 5.99% in the WT, adjusted p-val <0.001), immature α-like cells (Alpha; 10.95% in the KO versus 1.79% in the WT, adjusted p-val = 0.004), and ε-like cells (Epsilon; 14.63% in the KO versus 1.46% in the WT, adjusted p-val <0.001). Bipotent progenitors were present at equivalent proportions in both genotypes. The early *NEUROG3*+ population was expanded in NKX2.2KO cells (EP_early; 7.80% in the KO versus 4.10% in the WT, adjusted p-val = 0.032) consistent with accumulation at the endocrine progenitor stage. Although the late *NEUROG3*+ progenitor population was modestly reduced in the knockout (EP_late; 12.38% in the KO versus 17.60% in the WT), this difference was not significant (adjusted p-val = 0.445). Notably, the reduction in the late endocrine progenitors coincided with the expansion of the α-progenitor population. Collectively, these findings indicate that loss of NKX2.2 results in the accumulation of less mature endocrine progenitors and, among those cells that progress beyond this stage, a shift away from committed β- and δ-committed progenitor cell states towards α- and ε-committed progenitor populations.

Marker expression within each population revealed the same pattern of incomplete lineage specification (Figure 4E). NKX2.2KO cells assigned to the Beta_prog and Beta populations retained the progenitor-associated markers *HES6* and *PAX4* and failed to upregulate *NKX6-1* and *INS* to WT levels. Cells assigned to the Alpha_prog and Alpha populations expressed *ARX* and *ISL1* but showed reduced *ETV1* and *GCG*. Cells assigned to the Delta_prog and Delta populations contained fewer *HHEX* and *SST* expressing cells than the WT, while *GHRL* was broadly increased across the NKX2.2KO populations and the enterochromaffin markers *LMX1A*, *TPH1* and *SLC18A1* were reduced. Because clustering was performed on the merged dataset, NKX2.2KO cells assigned to a given cluster occupy the same transcriptional neighborhood as their WT counterparts but are not equivalent to them. Cluster identities should therefore be interpreted as positions along the differentiation trajectory rather than as fully established endocrine fates.

### The chromatin state of NKX2.2-bound elements predicts the direction of transcriptional change

To identify transcriptional changes that could reflect direct NKX2.2-dependent regulation, we reanalyzed a published NKX2.2 ChIP-seq dataset from human islets^42^. Peaks reproducible across the two donors were retained, giving 32,283 peaks that were then annotated using the human islet regulome^43^. The majority of peaks fell outside annotated regulatory elements (23,118, 71.6%), consistent with the known repressive role of NKX2.2^12,13^. Among the annotated sites, NKX2.2 occupied active enhancers (H3K4me1+/H3K27ac+; 2,603, 8.1%), inactive enhancers (H3K4me1+/H3K27ac-; 2,666, 8.3%), inactive open chromatin (H3K4me1-/H3K27ac-; 2,004, 6.2%), active promoters (982, 3.0%) and CTCF sites (910, 2.8%) (Figure 4F).

Interestingly, NKX2.2 occupancy alone did not predict transcriptional output. Across all cell populations, genes associated with an NKX2.2 peak were not significantly shifted in expression relative to unbound genes in any population (Wilcoxon adjusted p >0.5 for all partitions) however this is likely reflective of the biological differences between primary adult islets and stem cell-derived endocrine progenitors. Despite that difference, the chromatin context of the bound element was strongly predictive. Genes associated with NKX2.2-bound active enhancers or active promoters were preferentially downregulated in the NKX2.2 null cells, whereas genes associated with inactive enhancers were preferentially upregulated (Figure 4G; active enhancer versus inactive enhancer, p = 0.0065). Genes associated with both active and inactive enhancer binding sites also showed increased expression in the knockout.

To identify the biological processes most likely to be directly affected by NKX2.2 loss, we restricted the analysis to genes that were both NKX2.2-bound and differentially expressed and tested these gene sets for enrichment within each cell population (Figure 4H). Three coherent groups of terms emerged. The first comprised pancreatic and endocrine differentiation programs, including pancreas development, pancreatic β cell differentiation, pancreatic α cell differentiation, and enteroendocrine cell differentiation. The second comprised fate commitment processes, including cell fate commitment, pattern specification process and stem cell population maintenance. These were enriched across nearly all populations, including progenitor clusters. The third comprised WNT-related categories, including regulation of WNT and canonical WNT signaling. WNT terms were enriched across most of the endocrine populations in this analysis but did not emerge from the bulk pathway analysis, where gene expression changes in individual cell populations are averaged away. However, the critical WNT pathway gene *CTNNB1* was significantly increased at S5d4 (Figure 3G), suggesting NKX2.2 plays an important role in repressing the WNT pathway specifically in the endocrine progenitor population, where WNT inhibition has been described as a necessary step of endocrine progression^44^.

### NEUROG3 is not extinguished in NKX2.2KO cells

*NEUROG3* was upregulated in the NKX2.2KO cells and is associated with an NKX2.2-bound inactive enhancer, prompting us to directly examine its expression dynamics. In WT cultures, *NEUROG3* and *NKX2.2* were co-expressed within the same cells throughout the shared endocrine differentiation trajectory. Double-positive cells constituted the majority of the EP_early and EP_late populations and remained prevalent within the Beta_prog, Beta and Alpha_prog populations (Figure 5A, 5C). *NKX2.2*-only cells predominated in the Alpha population, corresponding to the stage at which *NEUROG3* is normally extinguished. Thus, the concurrent induction of *NEUROG3* and *NKX2.2* observed by qPCR (Figure 1B) reflects co-expression within individual cells rather than averaging across sequentially expressed cell populations. Co-immunofluorescence at S5d5 confirmed that NGN3 and NKX2.2 protein are both present in the WT endocrine progenitor population, and that NGN3 protein remains detectable in both NKX2.2KO clones (Figure 5B).

**Figure 5.**
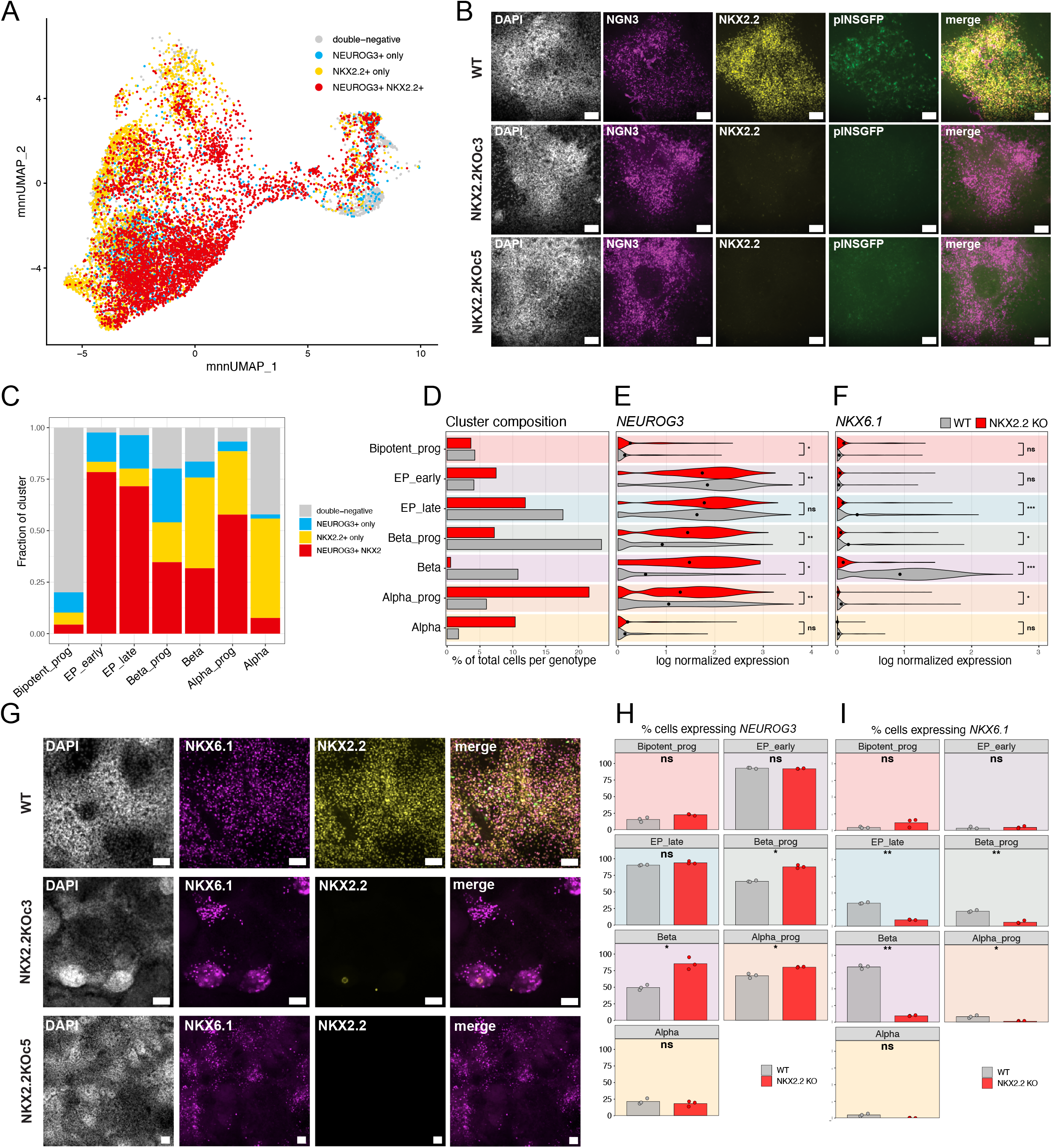
NEUROG3 and NKX2.2 are co-expressed, and NEUROG3 is not extinguished in NKX2.2KO cells. **(A)** UMAP of WT cells classified by *NEUROG3* and *NKX2-2* detection: double-negative, *NEUROG3*+ only, *NKX2-2*+ only, or double-positive. A cell was called positive at log normalized expression > 0.5. **(B)** Immunofluorescence for NGN3, NKX2.2 and pINSGFP with DAPI at S5d5 in WT and both NKX2.2 knockout clones. NGN3 and NKX2.2 protein are both present in WT endocrine progenitors, and NGN3 protein remains detectable in the knockout clones. Scale bars, 200 μm. **(C)** Fraction of WT cells in each population falling into the four classes in (A). Double-positive cells are the majority of EP_early and EP_late and remain prevalent in Beta_prog, Beta and Alpha_prog. *NKX2-2*-only cells predominate in the Alpha population, where *NEUROG3* is normally extinguished. **(D)** Selected population composition by genotype, as the percentage of each genotype’s cells assigned to each population. **(E)** *NEUROG3* expression per population by genotype, shown as violins of log normalized expression with the mean marked. Stars compare pseudobulk mean expression per differentiation by Welch’s two-tailed t-test, Benjamini-Hochberg adjusted across all 13 populations. *p < 0.05, **p < 0.01, ns, not significant. **(F)** *NKX6-1* expression per population by genotype, displayed and tested as in (E). *p < 0.05, ***p < 0.001, ns, not significant. **(G)** Immunofluorescence for NKX6.1 and NKX2.2 with DAPI at S5d4. Scale bars, 200 μm. Fields reused from Figure 3E. **(H)** Percentage of cells expressing *NEUROG3* (log normalized expression > 0) in each population, per independent differentiation. The increase in the knockout is significant in Beta_prog, Beta and Alpha_prog. **(I)** Percentage of cells expressing *NKX6-1* (log normalized expression > 0) in each population, per independent differentiation. *NKX6-1*-expressing cells are reduced in the knockout in EP_late, Beta_prog, Beta and Alpha_prog. For (H) and (I), bars are the mean of n = 3 differentiations per genotype with individual differentiations overlaid, compared by Welch’s two-tailed t-test, Benjamini-Hochberg adjusted across all 13 populations. *p < 0.05, **p < 0.01, ns, not significant.

In the NKX2.2KO cells, not only was *NEUROG3* expression significantly elevated (Supplemental Figure 3A-B), but it was also present in an expanded number of cells (Supplemental Figure 3C). Population-level analysis showed that NEUROG3 expression was not confined to the endocrine progenitor clusters (Supplemental Figure 3D) confirming that the committed cell populations were still in a progenitor state. Both the mean expression level (Figure 5E) and the proportion of *NEUROG3*-expressing cells (Figure 5H) were significantly increased in the Beta_prog, Beta and Alpha_prog populations, and mean expression was additionally decreased in EP_early populations, indicating there was both a delay in *NEUROG3* expression onset and persistent *NEUROG3* expression beyond the stage at which it is normally downregulated.

The same analysis applied to NKX6-1 produced the reciprocal result. In WT cultures, the majority of *NKX6-1*-expressing cells also expressed *NKX2.2* (Supplemental Figure 3F), and the two proteins were co-detected across the culture at S5d4 (Figure 5G). *NKX6-1*-expressing cells were far less frequent in the NKX2.2KO, falling from approximately 20% of cells in WT to approximately 3% (Supplemental Figure 3F). In combination with the identified occupancy of NKX2.2 in the active enhancer region of NKX6.1 (Figure 4I), this suggests that NKX2.2 is critical for the maintenance of NKX6.1 expression.

To examine the transcriptional effects of NKX2.2 loss along the differentiation trajectory, we reconstructed the α, β, δ and ε trajectories from the shared progenitor trunk using Slingshot (Figure 6A). Cell distributions along the trajectories differed markedly by genotype. WT cells populated the committed regions of the β and δ trajectories, whereas NKX2.2KO cells accumulated at intermediate pseudotime positions on the same trajectories, consistent with a failure to progress to fully differentiated states. There was also an observed shift of NKX2.2KO cells towards the α and ε trajectories (Figure 6B).

**Figure 6.**
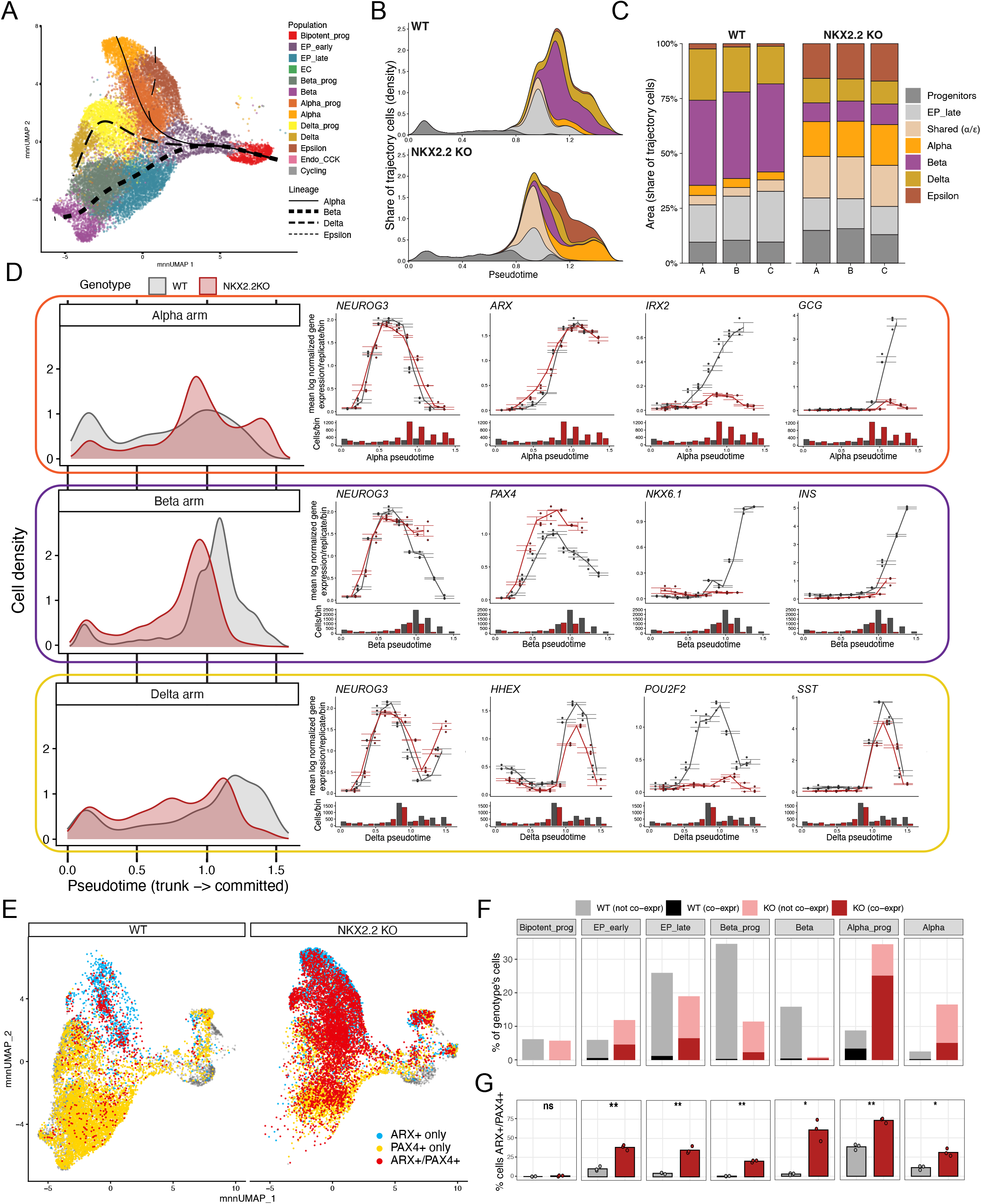
NKX2.2KO cells stall on the trajectory and fail to resolve a single lineage. **(A)** UMAP of the trajectory cells colored by annotated population, with the four fitted Slingshot lineages overlaid. Lineages were rooted at the bipotent progenitor population and terminate at the Alpha, Beta, Delta and Epsilon populations. **(B)** Density of trajectory cells along pseudotime for each genotype, stacked by lineage allocation. Cells on the α and ε curves prior to segregation are shown as shared (α/ε). Cells in the bipotent progenitor and EP_early populations are shown as progenitors. The NKX2.2KO density peaks earlier in pseudotime and does not recover at the committed end of the β and δ arms. **(C)** Share of trajectory cells in each lineage class, per independent differentiation. Unlike the population proportions in Figure 4D, which count cells against discrete cluster labels, this panel reports allocation between lineage arms and therefore captures cells that have not separated onto a single arm. The shared (α/ε) class rises from 4.5% in WT to 18.9% in the NKX2.2KO, the β share falls from 39.4% to 9.1%, and the progenitor share is higher in every knockout replicate (11.4 to 14.2%, compared with 9.2 to 10.2% in WT). **(D)** Left, density of cells along each of the α, β and δ trajectories by genotype. Right, mean expression of selected genes along the same trajectories, binned by pseudotime. Points are the mean of n = 3 independent differentiations per genotype with error bars showing standard error across differentiations, and the histograms beneath each panel give the number of cells per bin per genotype. *NEUROG3* fails to extinguish on the β and δ arms in the knockout, and *PAX4* remains elevated late on the β arm. *NKX6-1* and *INS* on the β arm, *IRX2* and *GCG* on the α arm, and *POU2F2* on the δ arm are induced only in WT, while *HHEX* and *SST* are induced in both genotypes. Each lineage was divided into 10 equal-width pseudotime bins. **(E)** UMAP split by genotype, with cells classified as *ARX*+ only, *PAX4*+ only, or double-positive, using log normalized expression thresholds of 1.0 for *ARX* and 0.5 for *PAX4*. Cells positive for neither are shown in grey. **(F)** Percentage of each genotype’s cells in each population, split by *ARX* and *PAX4* co-expression, using the thresholds in (E). **(G)** Percentage of cells co-expressing *ARX* and *PAX4* within each population, per independent differentiation. Co-expression increases in every endocrine population examined and is unchanged in bipotent progenitors. Welch’s two-tailed t-test on n = 3 differentiations per genotype, Benjamini-Hochberg adjusted. *p < 0.05, **p < 0.01, ns, not significant. Double-positive cells in (E) to (G) passed DoubletFinder filtering and were additionally checked against the doublet call.

Trajectory analysis also revealed differences between WT and NKX2.2KO cells that were not apparent from the relative abundance of the cell clusters alone (Figure 4D). Cluster analysis groups cells into discrete populations based on their transcriptional profiles and therefore provides a snapshot of the proportion of cells assigned to each cell identity. In contrast, trajectory analysis orders cells along inferred developmental lineages, allowing us to assess both how far cells have progressed through differentiation (pseudotime) and which endocrine lineage they are progressing toward. This analysis revealed two notable differences between WT and NKX2.2KO cells: first, the distribution of NKX2.2KO cells along pseudotime peaked earlier than that of WT cells, indicating that more NKX2.2KO cells accumulated at earlier stages of differentiation. NKX2.2KO cells were also underrepresented at the more differentiated ends of the β and δ cell lineage trajectories (Figure 6B). Second, approximately 19% of NKX2.2KO cells remained on the trajectory before the α and ε lineage paths diverged, compared with approximately 5% of WT cells (Shared α/ε, Figure 6C). This increase was accompanied by a decrease in the proportion of cells assigned to the β cell trajectory, from approximately 40% in WT to approximately 9% in NKX2.2KO cells. Consistent with these findings, the proportion of cells occupying the progenitor region of the trajectory was higher in every NKX2.2KO replicate (Figure 6C). Thus, whereas cluster proportions describe the relative abundance of cells with each transcriptional identity, trajectory analysis provides additional information about their position along the differentiation process and their progression toward specific endocrine lineages.

Consistent with the trajectory analysis, *NEUROG3* expression along the β cell lineage trajectory was induced normally in both genotypes but failed to be extinguished in NKX2.2KO cells (Figure 6D). Consequently, the fraction of *NEUROG3*+ cells in the late pseudotime window increased from 0.63 in WT to 0.87 in NKX2.2KO cells (p = 0.0007). A similar effect was observed along the δ trajectory (0.50 to 0.61, p = 0.006). In contrast, the difference along the α trajectory was not significant (p = 0.35) (Figure 6D).

The persistence of the progenitor program was accompanied by a failure to activate lineage-specific differentiation programs. Along the β cell trajectory, *NKX6-1* and *INS* increased sharply in WT cells but remained largely unchanged in the NKX2.2 KO cells, whereas *PAX4* expression peaked earlier and at higher levels. Along the α cell trajectory, *ARX* was induced similarly in both genotypes, however *IRX2* and *GCG* increased only in WT cells. Likewise, along the δ cell trajectory, *POU2F2* induction was restricted to WT cells, and expression of *HHEX* and *SST* remained lower in NKX2.2 KO cells (Figure 6D). Along the δ arm, *SST*, *HHEX* and *POU2F2* peaked at a pseudotime of approximately 1.2. Beyond this point, cells in the Delta population progressively lost *SST*, *HHEX*, and *POU2F2* and gained β cell genes (*INS, NKX6-1, MNX1, UCN3.* Supplemental Figure 4H-K). This occurred in both genotypes, consistent with instability of the δ state under β-directed culture conditions^45^.

Along the ε trajectory, NKX2.2KO cells were enriched at the committed end, consistent with the expansion of the Epsilon population (Figure 4D, Supplemental Figure 4A). *NEUROG3* was induced and extinguished with similar kinetics in both genotypes, although extinction was slightly delayed in NKX2.2KO cells (Supplemental Figure 4B). *GHRL* rose more slowly in NKX2.2KO cells but reached WT levels at the end of the trajectory (Supplemental Figure 4C). In contrast, the transcription factors that resolve endocrine fate were dysregulated. *PAX4* was induced earlier and peaked at higher levels in NKX2.2KO cells, and *ARX* was also induced earlier (Supplemental Figure 4D-E). *ACSL1* showed decreased expression in NKX2.2KO cells (Supplemental Figure 4F) indicating incomplete activation of late ε genes despite normal GHRL expression. *NEUROD1* was higher in NKX2.2KO cells from early pseudotime onward and remained elevated at the end of the trajectory (Supplemental Figure 4G). Together, these results indicate that NKX2.2KO cells reach the ε state in large numbers and acquire *GHRL* expression, but do so through a period of *PAX4* and *ARX* co-elevation and without full activation of the late ε program. This parallels the α, β and δ trajectories, where early lineage genes were induced but terminal programs were not.

This analysis revealed a common defect across all four endocrine lineages: genes associated with the endocrine progenitor state are not appropriately downregulated, while the transcriptional programs that define mature lineage identity fail to be fully established. For example, in the α cell lineage, *ARX* induction is preserved, suggesting that the α-directed cells initiate the appropriate lineage program, but fail to progress further down the lineage program, as evidenced by the absence of *IRX2* and GCG induction at later stages. This suggests there is a defect in progenitor-state resolution rather than lineage specification, consistent with the genetic positioning of *NKX2.2* downstream of *NEUROG3* in mouse endocrine development^15^.

### NKX2.2KO cells co-activate the mutually exclusive ARX and PAX4 programs

*ARX* and *PAX4* have been described to be cross-repressive transcription factors whose mutually exclusive expression has been proposed to establish the canonical lineage decision between α and β cells^37,46,47^. In the NKX2.2KO cells, however, both *ARX* and *PAX4* were upregulated. At the population level, simultaneous upregulation of these factors could reflect either co-expression within individual cells or expansion of distinct *ARX*- and *PAX4*-expressing populations. We therefore examined *ARX* and *PAX4* expression at the single-cell level to determine whether the two factors were expressed within the same cells.

*ARX*/*PAX4* double-positive cells were rare and spatially restricted in WT cultures but were widespread in the NKX2.2KO cultures (Figure 6E). Quantification across individual populations revealed significant increases in every endocrine compartment examined: EP_early (10% of the cells in the cluster to 38% in the KO), EP_late (7% to 30%), Beta_prog (2% to 20%), Beta (3% to 58%), Alpha_prog (38% to 72%) and Alpha (11% to 31%) (Figure 6F-G). In contrast, the fraction of double-positive cells in the bipotent progenitor population was unchanged, placing the onset of the defect at endocrine commitment rather than at an earlier developmental stage. All double-positive cells passed DoubletFinder filtering and were classified as singlets.

Together with the trajectory analyses, these findings define the altered developmental state of NKX2.2KO endocrine cells at the endocrine progenitor stage. *NEUROG3* expression persists beyond its normal developmental window, the normally mutually exclusive *ARX*-*PAX4* lineage decision fails to resolve appropriately, and the terminal transcriptional programs of the α, β, δ and ε lineages remain incompletely established. The hormone-poor, polyhormonal and hormone-negative cells observed at the end of the differentiation (Figure 2C-F) are consistent with a failure to fully resolve endocrine progenitor identity and establish mature, lineage-specific transcriptional programs.

## DISCUSSION

The purpose of this study was to determine the precise molecular role of NKX2.2 during human pancreatic islet development. Homozygous hESC lines deleted for *NKX2.2* were guided through a β cell differentiation protocol and analyzed at the endocrine progenitor stage and at the end of the differentiation, allowing us to determine the earliest functions of NKX2.2. Specification of the endocrine lineage did not require NKX2.2: *NEUROG3* was induced normally, *CHGA*+ cells were generated at close to wild-type frequency, and differentiation into the endocrine lineages occurred appropriately. However, upon endocrine cell specification, the NKX2.2KO cell lines were impaired in their ability to extinguish *NEUROG3* and *ARX* and *PAX4* became co-activated within the same cells. Furthermore, the terminal specifiers of the α, β and δ lineages were not induced, resulting in few insulin-producing cells, an excess of somatostatin and ghrelin expressing cells, polyhormonal cells, and a large hormone-negative population. These results indicate that NKX2.2 is required for the proper, mono-hormonal resolution of endocrine fate in human pancreas development.

Although the endpoint phenotypes differ between mouse and human, this can partly be explained by the differentiation protocol. Mice lacking *Nkx2.2* have no β cells, few α cells and a large expansion of ghrelin-expressing ε cells^10,11^. The NKX2.2KO cultures lose most *INS*+ cells and gain *GHRL*+ cells, although the gain is smaller than in mice, and *GCG*+ cells are rare in both genotypes. The protocol used here is heavily biased toward the β lineage and produces few α cells even in the wild type context, so it is not surprising there was not a measurable decrease in α cell numbers. It is also possible that the increase in SST+ cells is unique to human development. The human patient data are consistent with what we observe in the compartments that were measured, namely absent C-peptide at birth and elevated serum ghrelin (Flanagan, Cell Metab, 2014).

The increase in somatostatin at the end stage of differentiation was inconsistent with its expression at the endocrine progenitor stage. At S5d4, the δ specifier *POU2F2* is among the most strongly downregulated genes in the NKX2.2KO cell lines, while *HHEX* is modestly reduced. Consistent with this, *HHEX*, *POU2F2* and *SST* all show impaired induction along the δ trajectory. Yet by S6d14, SST+ cells are increased by an order of magnitude and *HHEX* transcript is correspondingly elevated approximately two-fold. These findings can be reconciled if the late-emerging SST+ cells lack a fully specified δ cell identity, as is suggested by the near absence of *POU2F2* transcript along the δ pseudotime trajectory. In this interpretation, somatostatin expression occurs despite the failure to establish the δ cell transcriptional program during progenitor differentiation, reflecting hormone expression in cells with unresolved endocrine identities rather than a true expansion of the δ lineage. The ghrelin-expressing population presents a different picture. Single-cell analysis demonstrates an expansion of the ε-cell compartment in the knockout, evident both in cluster abundance (Figure 4D) and in the proportion of trajectory cells assigned to the ε branch (Figure 6C). This finding is consistent with previous reports of NKX2.2 loss in humans and mice.

Persistent *NEUROG3* expression is the most striking feature of the NKX2.2 knockout state. Under normal conditions, *NEUROG3* is expressed transiently, and the developmental competence conferred by that pulse of expression changes over time^9^. Thus, both the onset and termination of *NEUROG3* expression are important for endocrine differentiation. In NKX2.2KO cells, *NEUROG3* expression initiates normally, but fails to terminate. *NEUROG3* remains detectable in cells progressing along the β and δ trajectories, and is elevated even in populations that have been assigned to committed endocrine identities. NKX2.2 occupies a single regulatory element at *NEUROG3*, approximately 3 kb from the transcription start site, that is classified as an inactive-enhancer, consistent with increased gene expression following NKX2.2 loss. Together, these findings support a model in which NKX2.2 directly represses *NEUROG3* to ensure its timely extinction. Alternatively, increased *CTNNB1* expression in the NKX2.2KO could suggest an indirect mechanism through which NKX2.2 inhibits *CTNNB1* expression to allow for endocrine progression. NKX2.2 occupies several distal sites upstream of *CTNNB1* (data not shown) although these sites were unannotated in the Human Islet Regulome, so we present this only as a possible alternative mechanism. The absence of detectable NKX2.2 binding at *HES1* argues against direct regulation of *HES1* as a mechanism underlying this effect, although NOTCH-dependent or other indirect mechanisms cannot be excluded. Ultimately, the proposed direct repression of NEUROG3 remains correlative and will require functional validation through deletion or mutation of the bound enhancer.

Co-expression of *ARX* and *PAX4* within individual cells provides a potential transcriptional explanation for the emergence of polyhormonal cells. *ARX* and *PAX4* are mutually repressive, and resolution of their expression is central to the α versus β lineage decision^37,47^. In the NKX2.2 knockout, a substantial fraction of cells across all endocrine populations express both factors, including cells already assigned to the β and α cell lineages. Polyhormonal cells are a persistent and undesirable outcome of directed endocrine differentiation protocols, and contribute to the failure of appropriate hormone regulation. Our findings instead suggest a more fundamental defect in lineage specification: cells enter the endocrine compartment without resolving the competing transcriptional programs that determine their identities. If this mechanism extends beyond *NKX2.2* loss, *ARX* and *PAX4* co-expression should also occur during wild-type differentiations at frequencies related to the emergence of polyhormonal cells. Consistent with this possibility, double-positive cells compose only 6.22% of the wild-type endocrine populations but constitute 44.7% of the NKX2.2KO populations (Supplemental Figure 4L). In both cell lines, the majority of co-expressing cells were present in the Alpha_prog cluster (Figure 6F-G), potentially reflecting the β cell bias of the differentiation protocol. This observation is compatible with the proposed mechanism but does not establish a causal relationship between *ARX*/*PAX4* co-expression and polyhormonal identity.

Trajectory analysis provides an independent, annotation-free perspective on this failure of lineage resolution. Cells that remain positioned between adjacent lineage branches and cells that co-express the transcriptional regulators of competing fates may represent complementary manifestations of the same underlying defect: an inability to resolve alternative endocrine identities. Thus, the trajectory and transcriptional analyses converge on a model in which NKX2.2 is required not simply for appropriate hormone expression, but for the resolution and stabilization of endocrine cell fate.

The hormone-negative population present at the end of the differentiation provides particularly strong support for the lineage-resolution model. At S6d14, approximately 25–33% of NKX2.2KO cells express none of the three major endocrine hormones (*INS*, *GCG*, or *SST*), compared with only a small fraction of wild-type cells. These cells appear to have differentiated into endocrine lineages, as the proportion of *CHGA*+ cells is largely preserved at S6d1, but subsequently fail to express a hormone. This distinction is important: a defect in endocrine specification would be expected to reduce entry into the endocrine compartment, whereas a defect in lineage resolution would allow cells to enter, but leave them without a defined endocrine identity. The latter is more consistent with our observations. Importantly, this hormone-negative population can be overlooked when differentiation efficiency is assessed only among hormone-expressing cells. We therefore suggest that the proportion of hormone-negative cells should be routinely included in the phenotypic assessment of directed endocrine differentiation.

Interestingly, the presence of an NKX2.2 peak near a gene, by itself, did not predict whether that gene was differentially expressed in the knockout or the direction of its response. In contrast, the chromatin state of the bound regulatory element was informative: occupancy at active enhancers and promoters was associated with reduced expression following NKX2.2 loss, whereas occupancy at inactive enhancers was associated with increased expression. An important limitation, however, is that the ChIP data were obtained from adult human islets, whereas differential expression was measured in endocrine progenitors. The chromatin annotations therefore reflect a later developmental state and do not necessarily represent the regulatory state of the same elements at S5d4.

Loss of *NKX2.2* also markedly reduces the enterochromaffin population, a common off-target product of β cell differentiation protocols^26^. Key components of the enterochromaffin transcriptional program, including *LMX1A*, *TPH1* and *SLC18A1*, are strongly downregulated in the knockout at both the progenitor and terminal stages. *FEV* expression is similarly reduced, consistent with NKX2.2 acting upstream of the enterochromaffin differentiation program. These findings indicate that NKX2.2 is required for the formation of this population. Although reducing enterochromaffin population is a desirable goal for differentiation protocol development, its loss is accompanied by severe disruption of the β cell lineage. Thus, NKX2.2 inhibition does not provide a selective means of eliminating enterochromaffin cells, but the finding may help identify regulatory mechanisms that could eventually be targeted more specifically.

Taken together, these findings identify NKX2.2 as a critical regulator of the transition from endocrine specification to stable endocrine identities in human pancreatic differentiation. This distinction has implications beyond NKX2.2. End-stage hormone expression alone cannot distinguish a defect in endocrine specification from a failure to resolve and maintain a particular endocrine fate, yet these defects may require different strategies to correct. Current directed differentiation protocols can efficiently generate endocrine cells, but the resulting populations remain heterogeneous and incompletely mature compared with primary human islets^48^. Our findings suggest that a key remaining challenge is not simply generating endocrine cells, but ensuring that they extinguish progenitor programs and resolve competing lineage identities. Understanding the transcriptional mechanisms that govern this transition may therefore provide a productive route toward improving the fidelity and maturity of stem-cell-derived endocrine populations.

## LIMITATIONS OF THE STUDY

This study used a single parental hESC line (MEL-1 INS^GFP/w^, male) and one directed differentiation protocol, so the endpoint phenotype may differ in other genetic backgrounds or under protocols with a different lineage balance. However, we addressed clonal variation by analyzing two independent NKX2.2KO clones, which gave concordant phenotypes. These phenotypes also matched the available mouse and patient data, namely loss of insulin and elevated ghrelin. The protocol is strongly biased toward the β lineage and produces few α cells even in WT cultures, so the effect of NKX2.2 loss on α cell abundance is likely underestimated. To account for this, we assessed the α lineage by progression along its trajectory, using *IRX2* and *GCG* induction, rather than by α cell number. In addition, S6d14 cells remain immature relative to primary islets. We therefore located the primary defect at the progenitor stage, and at S6d14 we quantified hormone-negative cells as well as hormone-expressing cells. We did not measure hormone secretion or test function after transplantation, so the functional consequences of the hormone-poor and hormone-negative populations remain undefined.

The single-cell data come from one timepoint, S5d4. Pseudotime orders cells by transcriptional similarity and does not measure developmental time or establish lineage relationships. The persistence of *NEUROG3* along the β and δ arms is therefore inferred from how cells are ordered. However, the same result was seen without pseudotime: the fraction of *NEUROG3*+ cells was increased in the Beta_prog, Beta and Alpha_prog clusters. Trajectories were fitted jointly to both genotypes with a fixed δ branch point, and NKX2.2KO cells may not follow the same paths as WT cells. We checked the fixed branch point against an unconstrained fit, and the δ results did not change. Several clusters were small in one genotype, including Beta and Delta in the knockout and Epsilon in WT, which limits the precision of estimates for those populations. To limit the effect of small populations, we treated the independent differentiation as the unit of replication for all statistics. We also excluded pseudobulk profiles built from fewer than 10 cells.

The link between the S5d4 transcriptional state and the S6d14 hormone phenotype is inferred. We did not perform single-cell profiling or lineage tracing at S6d14. We therefore cannot show that *ARX*/*PAX4* co-expressing cells give rise to polyhormonal or hormone-negative cells, or that the late SST+ cells lack δ identity. We partly addressed this with bulk RNA-seq of the same clones at both timepoints. This showed that *SST* and *HHEX* elevation arises after S5d4 rather than at the onset of the defect. Also, all *ARX*/*PAX4* double-positive cells passed doublet filtering, so co-expression reflects single cells rather than captured pairs.

Finally, NKX2.2 occupancy came from published ChIP-seq of adult human islets, and chromatin state was assigned from an adult islet regulome. Binding and chromatin state at S5d4 may differ, and ChIP on bulk islets cannot assign binding to a specific endocrine cell type. However, we retained only peaks reproducible across both donors. We also checked regulome assignment against a known MAFA enhancer as a positive control. Rather than relying on occupancy alone, which did not predict expression change, we partitioned bound genes by chromatin state. Peaks were assigned to the nearest gene, which may not be the gene they regulate. For this reason we restricted the enrichment analysis to genes that were both bound and differentially expressed. We also used the published peak calls rather than reprocessing the reads. The proposed direct regulation of *NEUROG3*, *NKX6-1* and other targets therefore remains correlative. Testing it will require ChIP in differentiating cells and perturbation of the bound elements.

## RESOURCE AVAILABILITY

### Lead contact

Further information and requests for resources and reagents should be directed to and will be fulfilled by the lead contact, Lori Sussel

### Materials availability

The NKX2.2KO MEL-1 INS^GFP/w^ lines generated in this study are available from the lead contact.

### Data and code availability

- Bulk and single cell RNA sequencing data have been deposited at GEO and are publicly available as of the date of publication; accession numbers are listed in the key resources table. Previously published NKX2.2 ChIP-seq data are available at ArrayExpress under E-MTAB-1919.
- All original code used for the analyses in this paper is available at https://github.com/CUAnschutzBDC/snakemake_pipelines.
- Any additional information required to reanalyze the data reported in this paper is available from the lead contact upon request.

## ACKNOWLEDGMENTS

This work was supported by grants from the National Institutes of Health P30DK116073, R01 DK082590, and U01 DK127505 to LS. The authors acknowledge Dr. Holger Russ for his early guidance in establishing hESC culture and for sharing the MEL-1 INS^GFP/w^ cell line. This work utilized the Alpine High-Performance Computing resource at the University of Colorado Boulder. Alpine is jointly funded by the University of Colorado Boulder, the University of Colorado Anschutz, Colorado State University, and the National Science Foundation (award 2201538). We would also like to acknowledge the Barbara Davis Center Flow Cytometry core facility, and the University of Colorado Diabetes Research Center Disease Modeling Core, both supported by a grant from the National Institute of Diabetes and Digestive and Kidney Diseases P30-DK116073. Finally, we thank the University of Colorado Cancer Center Genomics Shared Resource (P30CA046934) for providing library preparation and sequencing services

## AUTHOR CONTRIBUTIONS

Conceptualization, CS and LS; experimental design and performance, CS, LS, MXR, FMD; cell line generation, PSM; data analysis, CS, CJH and KLW; writing—original draft, CS and LS; writing—review & editing, CS, LS, MXR, FMD, KLW; funding acquisition, LS.

## DECLARATION OF GENERATIVE AI AND AI-ASSISTED TECHNOLOGIES IN THE WRITING PROCESS

During the preparation of this work, the author(s) used Claude (Anthropic, claude.ai) in order to assist with grammar and wording, code planning/review, and to assist in drafting the methods section based on author-provided code and analytical decisions. After using this tool or service, the author(s) reviewed and edited the content as needed and take(s) full responsibility for the content of the publication.

## SUPPLEMENTAL INFORMATION

Document S1. Supplemental Figures S1-S6, Supplemental table 1

---

**Supplemental Figure 1.**
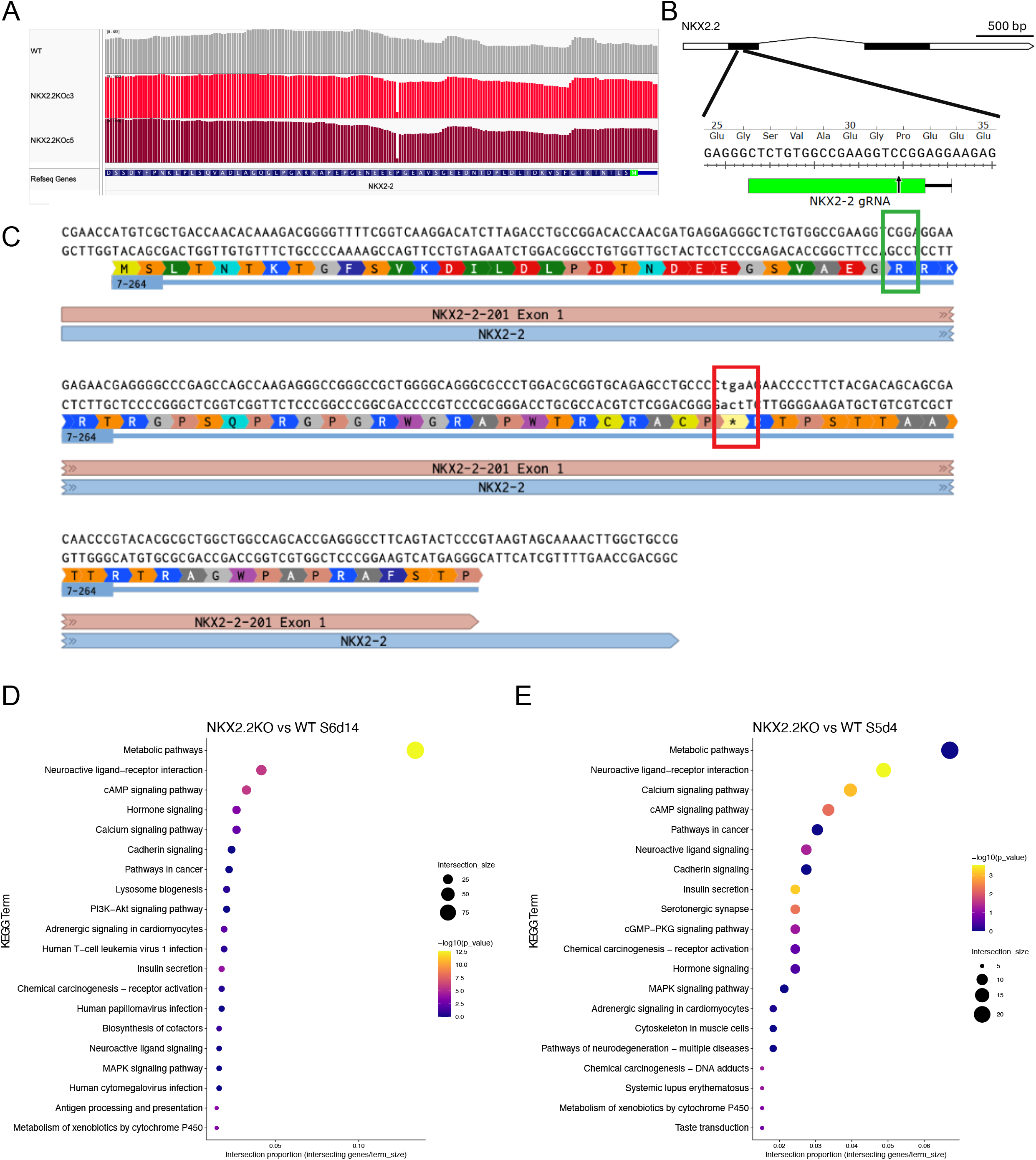
Generation of NKX2.2KO hESC lines and pathway enrichment of genes differentially expressed in NKX2.2KO cells. **(A)** IGV tracks of bulk RNA-seq reads at the NKX2.2 locus in WT cells and the two NKX2.2KO clones (c3 and c5), generated in the MEL1 INS^GFP/w^ line. Both clones carry the same single base pair deletion in exon 1. **(B)** Schematic of the NKX2.2 locus, with the gRNA target in exon 1 shown against codons 25–35. The arrow marks the cut site. Scale bar, 500 bp. **(C)** Sequence of NKX2.2 exon 1 in the NKX2.2KO allele with amino acid translation. The green box marks the cut site, where loss of a single cytosine shifts the reading frame. The red box marks the resulting premature stop codon. **(D, E)** KEGG overrepresentation analysis of genes differentially expressed between NKX2.2KO and WT cells by bulk RNA-seq at S6d14 **(D)** and S5d4 **(E)**. Differentially expressed genes were called by DESeq2 at adjusted p < 0.05 with no fold change threshold, combining both NKX2.2KO clones against WT. Enrichment was tested with g:Profiler. The 20 terms with the highest intersection proportion (differentially expressed genes in the term divided by term size) are shown, ranked by that proportion. Point size shows the number of differentially expressed genes in each term, and color shows −log10 adjusted p (g:SCS). n = 3 independent differentiations per cell line.

**Supplemental Figure 2.**
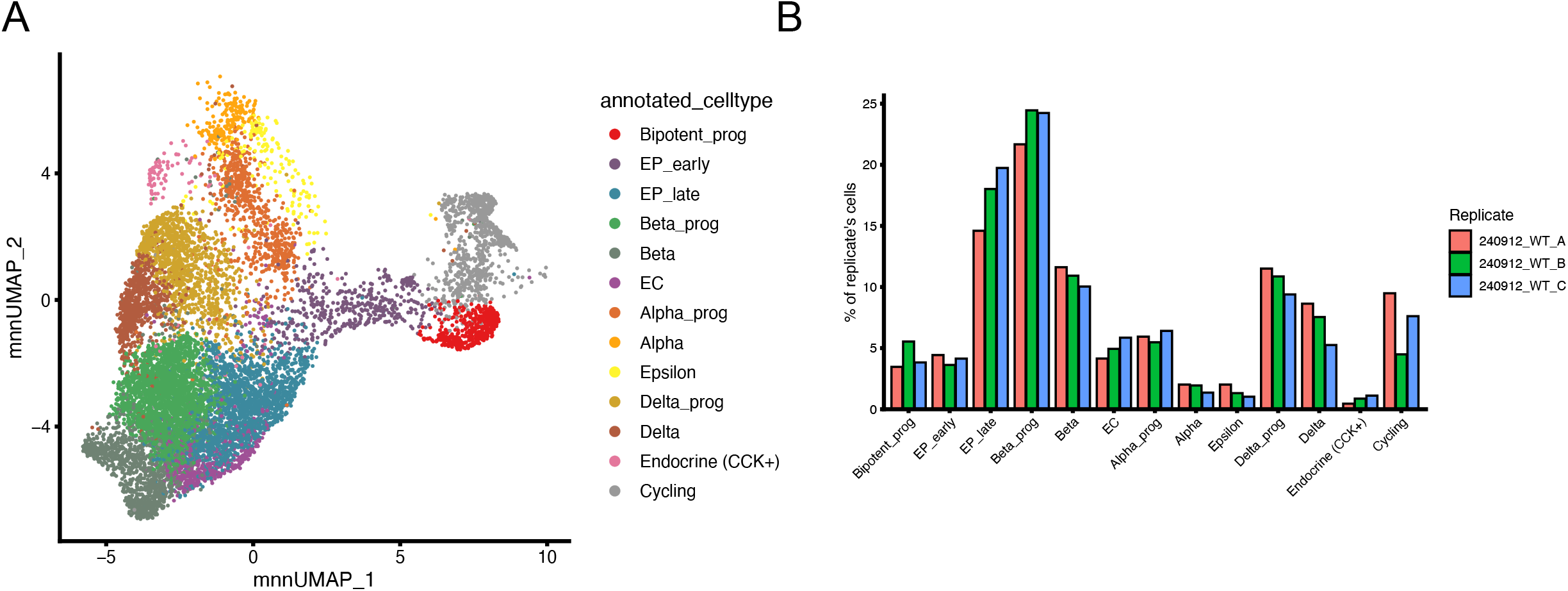
Population annotation and composition of WT cells. **(A)** UMAP of WT cells from the joint WT and NKX2.2KO embedding, colored by annotated population. Populations were named from marker expression in WT cells only (WT vote), and the same labels were applied to NKX2.2KO cells in each cluster. **(B)** Percentage of each WT replicate’s cells assigned to each population. Each bar is one independent differentiation (240912_WT_A, B and C).

**Supplemental Figure 3.**
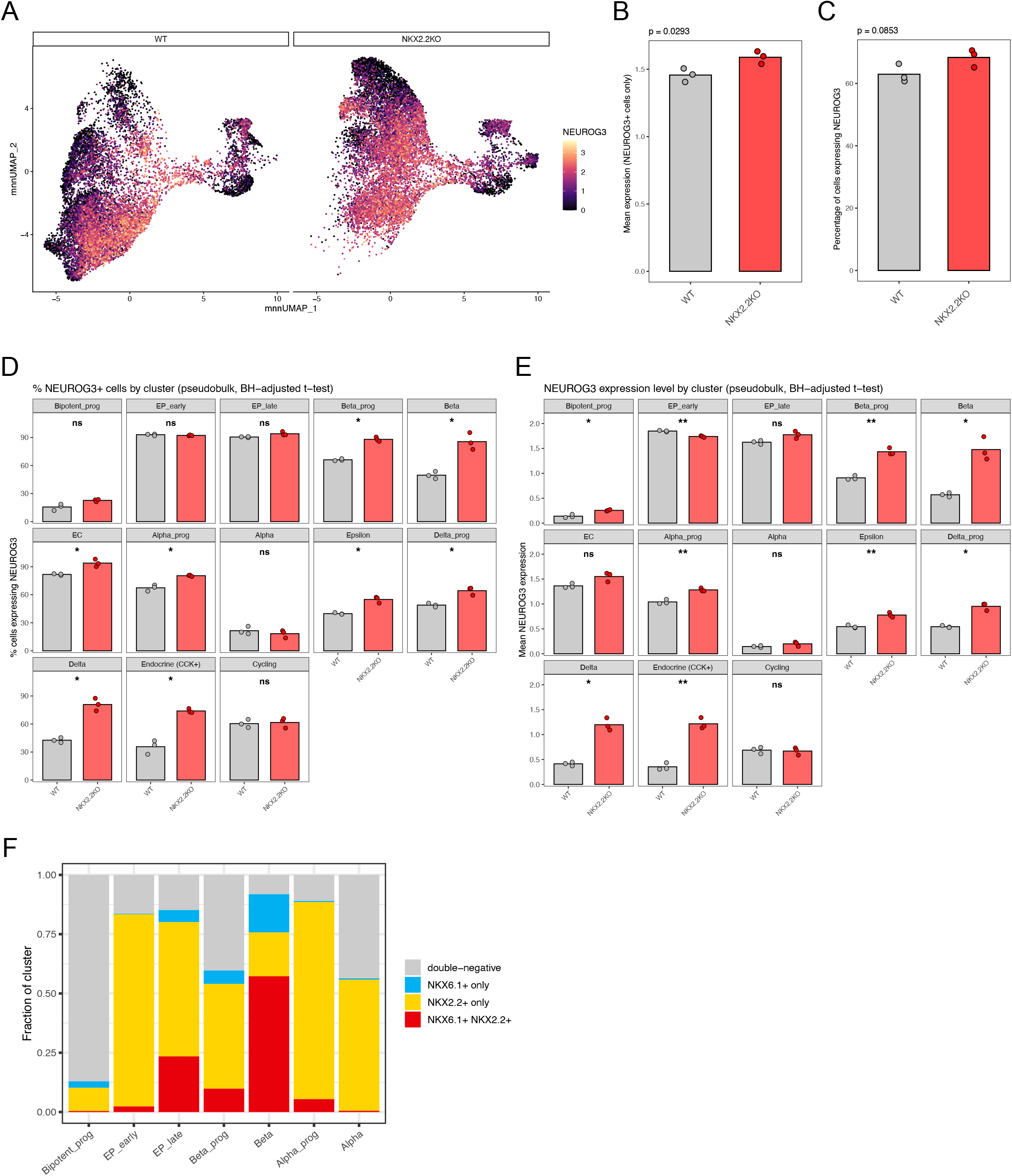
NEUROG3 expression and NKX6.1/NKX2.2 co-expression across populations. **(A)** *NEUROG3* expression (log-normalized) on the joint UMAP, split by genotype. **(B)** Mean *NEUROG3* expression in *NEUROG3*+ cells and **(C)** percentage of all cells expressing *NEUROG3*, per independent differentiation across the whole dataset. **(D)** Percentage of *NEUROG3*+ cells and (E) mean *NEUROG3* expression in each population, calculated per independent differentiation (pseudobulk). In B–E, a cell was scored *NEUROG3*+ if any transcript was detected. **(F)** Fraction of WT cells in each population that were *NKX6.1*+ *NKX2.2*+, *NKX2.2*+ only, *NKX6.1*+ only or double-negative, with a cell scored positive at log-normalized expression above 0.5. Bars show the mean of three independent differentiations per genotype, with points for each differentiation. Genotypes were compared by two-tailed Welch’s t-test. In D and E, p values were adjusted by Benjamini-Hochberg across the 13 populations. ns, not significant; *p < 0.05; **p < 0.01.

**Supplemental Figure 4.**
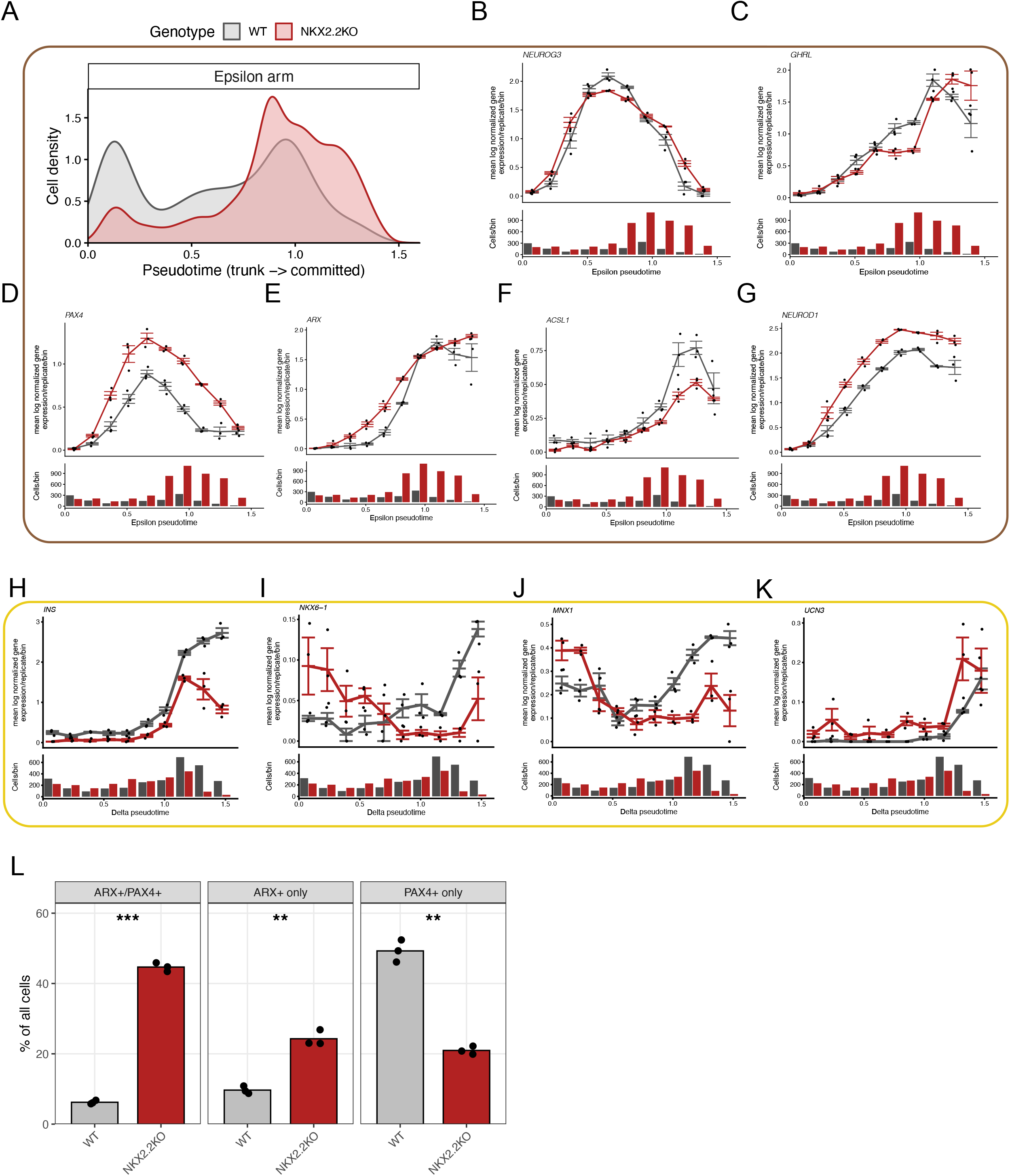
The ε trajectory, β cell genes along the δ trajectory, and ARX/PAX4 co-expression. **(A)** Density of WT and NKX2.2KO cells along the ε trajectory of the joint Slingshot fit. A cell was assigned to a trajectory if its curve weight was 0.5 or greater. **(B–G)** Expression of *NEUROG3* **(B)**, *GHRL* **(C)**, *PAX4* **(D)**, *ARX* **(E)**, *ACSL1* **(F)** and *NEUROD1* **(G)** along ε pseudotime. **(H–K)** Expression of the β cell genes *INS* **(H)**, *NKX6-1* **(I)**, *MNX1* **(J)** and *UCN3* **(K)** along the full δ pseudotime. In **(B–K)**, points are mean log-normalized expression per differentiation in each equal-width bin, and lines and bars show the mean ± SEM of three independent differentiations per genotype. Cells per bin are shown beneath each panel. **(L)** Percentage of all cells that were *ARX*+/*PAX4*+, *ARX*+ only or *PAX4*+ only, per independent differentiation across the whole dataset. A cell was scored positive at log-normalized expression above 0.5. Bars show the mean, with points for each differentiation. Genotypes were compared by two-tailed Welch’s t-test with Benjamini-Hochberg correction across the three states. **p < 0.01; ***p < 0.001.

**Supplemental Figure 5.**
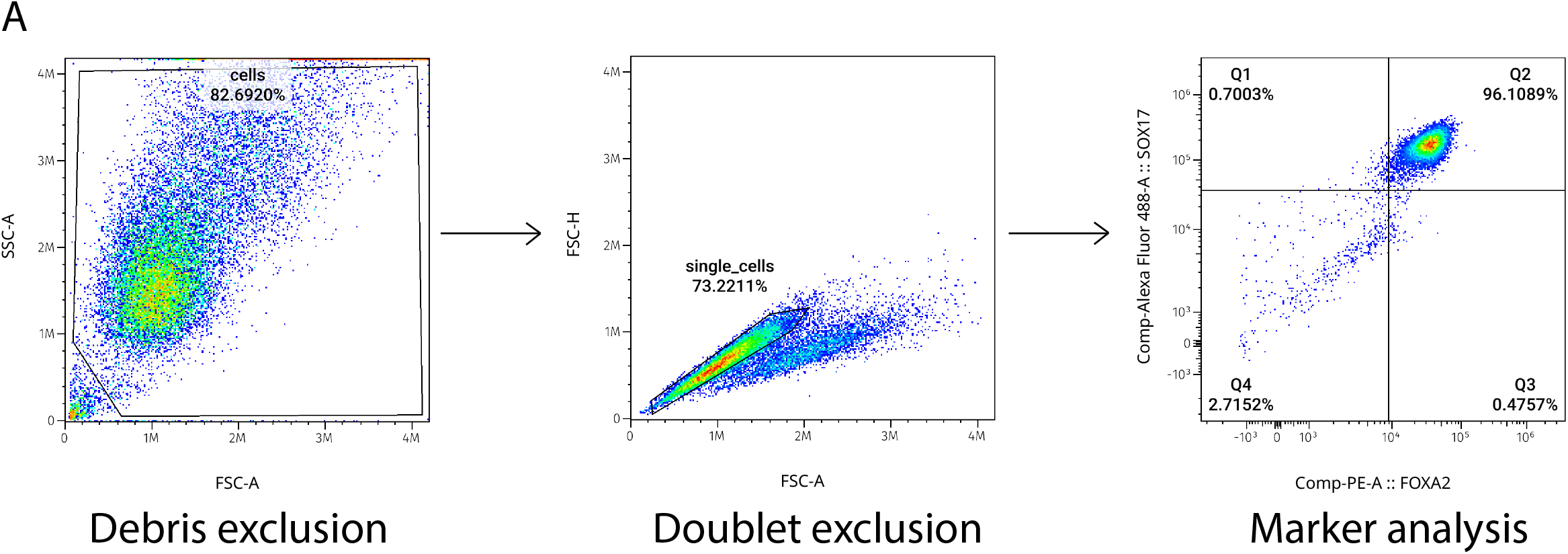
Flow cytometry gating strategy. **(A)** Representative gating for all flow cytometry experiments. Debris was excluded on forward and side scatter area (FSC-A vs SSC-A), and doublets were excluded on forward scatter area versus height (FSC-A vs FSC-H). Single cells were then analyzed for panel specific marker expression.

**Supplemental Figure 6.**
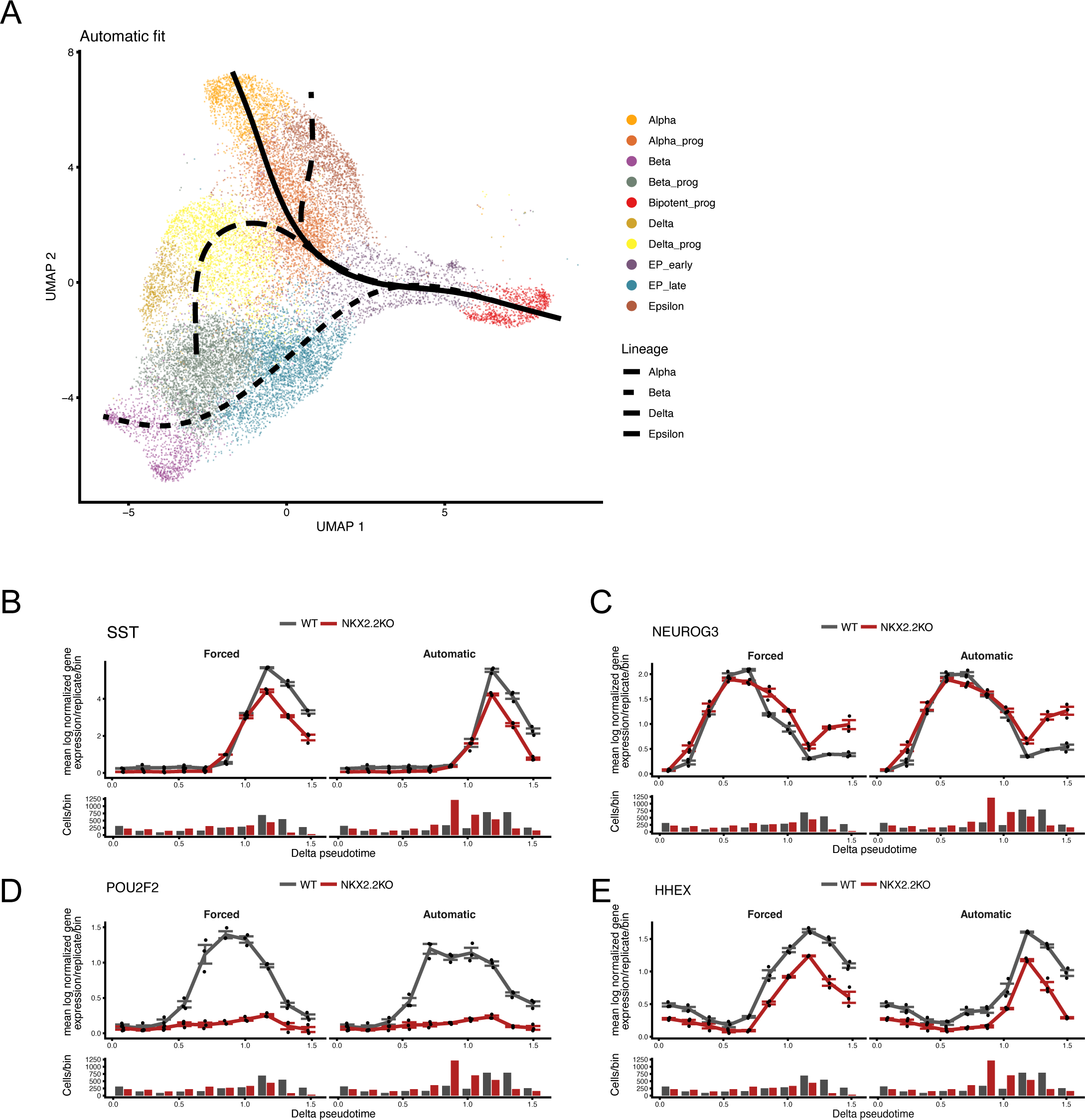
The δ trajectory under an unconstrained Slingshot tree. **(A)** UMAP of trajectory cells with curves from a joint Slingshot fit in which the tree was chosen automatically, using the same start (Bipotent_prog) and endpoints (Alpha, Beta, Delta and Epsilon) as the main fit. The α, β and ε lineages match the fixed tree, and the δ lineage passes through Alpha_prog. **(B–E)** Expression of *SST* **(B)**, *NEUROG3* **(C)**, *POU2F2* **(D)** and *HHEX* **(E)** along δ pseudotime in the fixed (Forced, left) and unconstrained (Automatic, right) fits. Points are mean log-normalized expression per differentiation in each equal-width bin, and lines and bars show the mean ± SEM of three independent differentiations per genotype. Cells per bin are shown beneath each panel.

**Supplemental Table 1.** Primary antibody dilutions.

| Antibody | Source | Conjugate | Application | Dilution used |
|---|---|---|---|---|
| Mouse anti-Sox2 | BD Biosciences | Alexa Fluor 647 | Flow cytometry | 1:100 |
| Mouse anti-Human TRA-1-60 | BD Biosciences | PE | Flow cytometry | 1:200 |
| Mouse anti-PDX1 | BD Biosciences | PE | Flow cytometry | 1:20 |
| Mouse anti-NKX6.1 | BD Biosciences | Alexa Fluor 647 | Flow cytometry | 1:50 |
| Rabbit anti-Insulin | Cell Signaling Technologies | Alexa Fluor 488 | Flow cytometry | 1:50 |
| Mouse anti-Chromogranin A | Novus Biologicals | none | Flow cytometry | 1:50 |
| Mouse anti-NKX2.2 | DSHB | none | Immunofluorescence | 1:50 |
| Sheep anti-NEUROG3 | R&D Systems | none | Immunofluorescence | 1:300 |
| Rat anti-C-peptide | DSHB | none | Immunofluorescence | 1:1,000 |
| Goat anti-PDX1 | R&D Systems | none | Immunofluorescence | 1:200 |
| Mouse anti-NKX6.1 | DSHB | none | Immunofluorescence | 1:200 |
| Rabbit anti-Somatostatin | Phoenix Pharmaceuticals | none | Immunofluorescence | 1:200 |
| Rabbit anti-Ghrelin | Phoenix Pharmaceuticals | none | Immunofluorescence | 1:200 |
| Goat anti-NKX6.1 | R&D Systems | none | Immunofluorescence | 1:500 |

## STAR★METHODS

### KEY RESOURCES TABLE

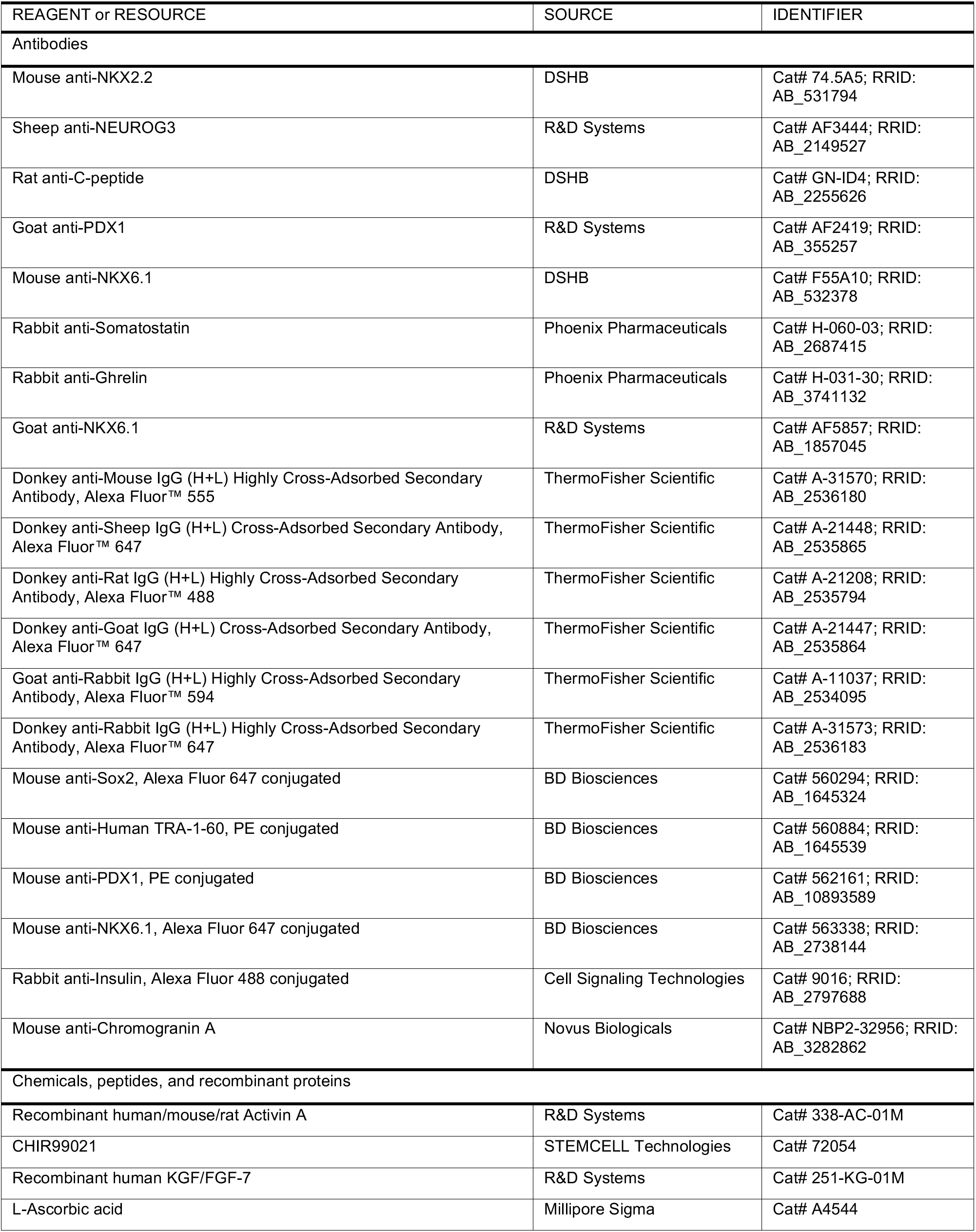
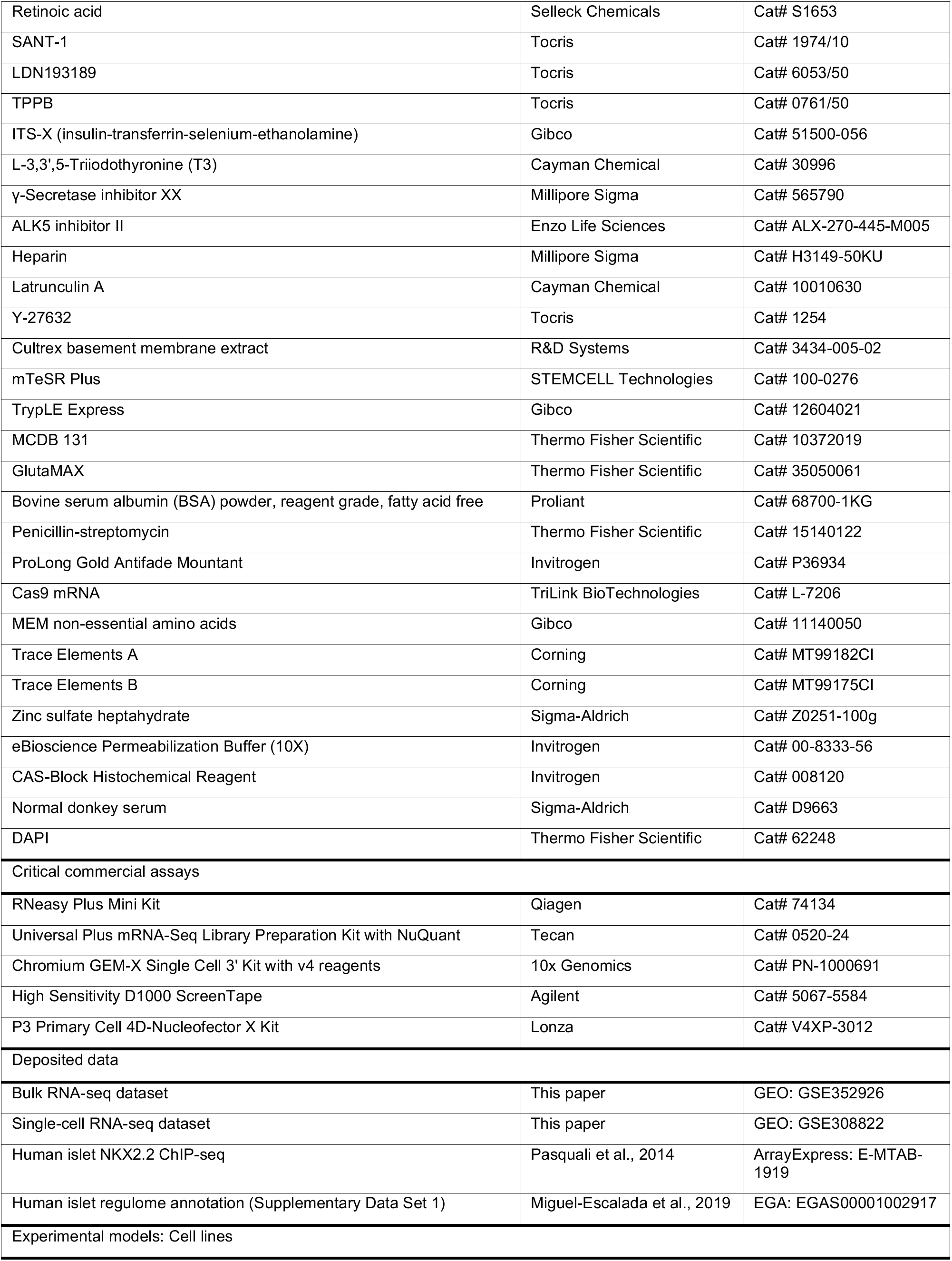
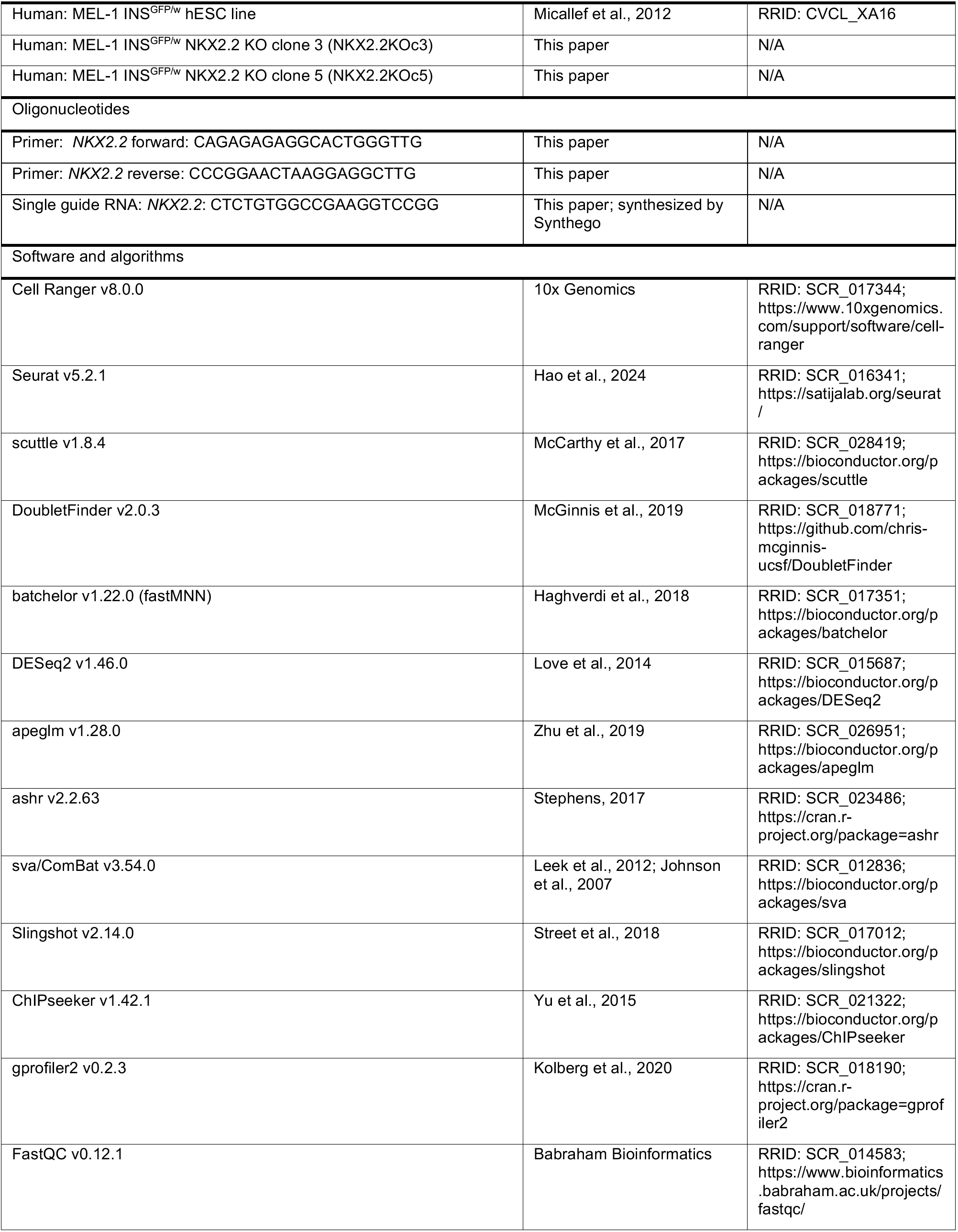
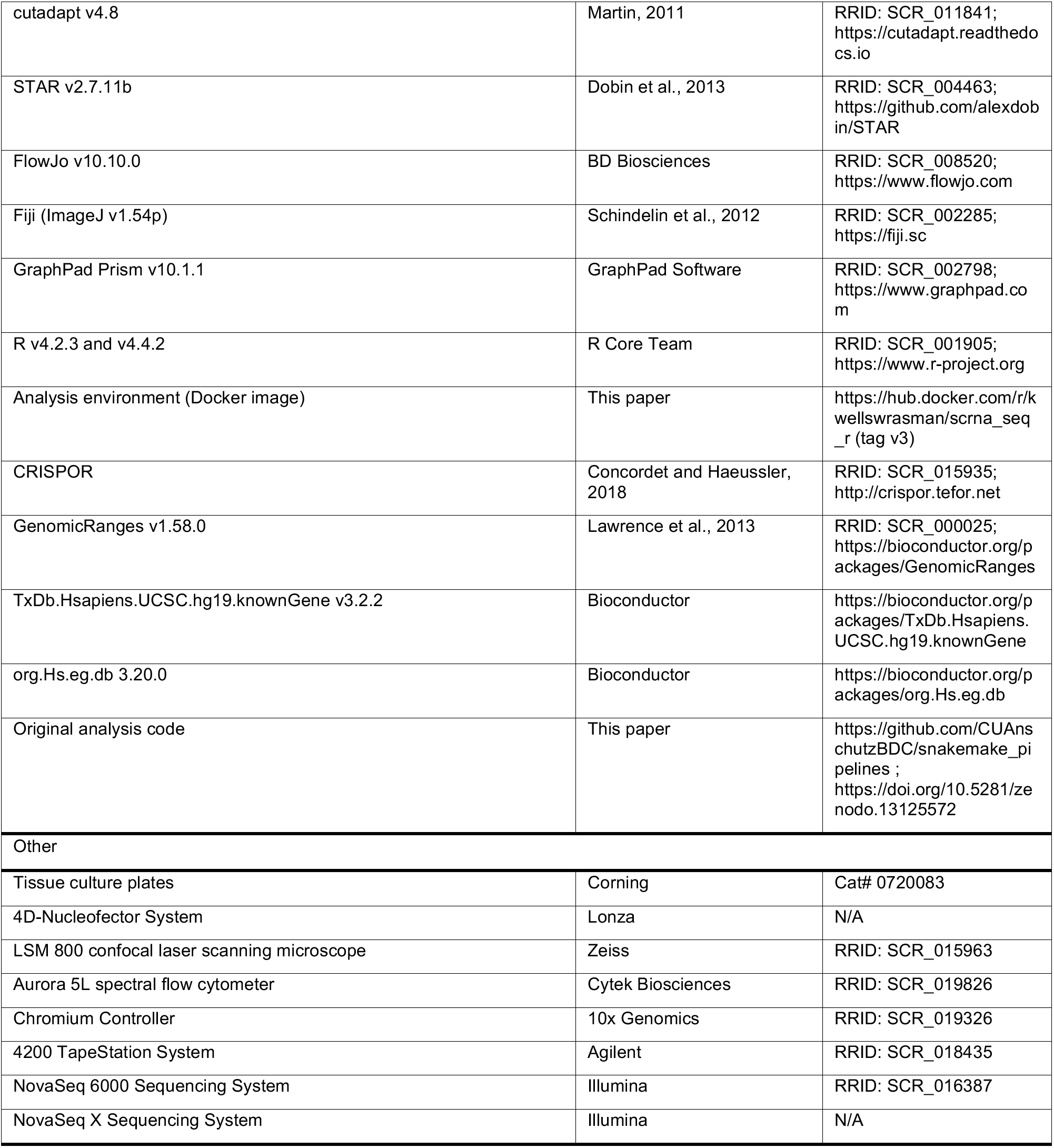

### EXPERIMENTAL MODEL AND STUDY PARTICIPANT DETAILS

MEL-1 INS^GFP/w^ (RRID:CVCL_XA16) is a male human embryonic stem cell line carrying GFP under the control of the INS promoter, and was generously provided by Dr. Holger Russ. NKX2.2KOc3 and NKX2.2KOc5 were derived from this parental line as described below, and chromosomal integrity of the WT cell line and both NKX2.2KO clones was confirmed by karyotyping performed by the University of Colorado Cancer Center Molecular Pathology Shared Resource. Mycoplasma testing was performed by the Barbara Davis Center Bioresources Core Molecular Biology Unit and confirmed negative for all cell lines. No human participants and no animals were used in this study.

#### Generation of NKX2.2KO lines

NKX2.2KO lines were generated at the Disease Modeling Core, Barbara Davis Center for Diabetes. The NKX2-2 guide RNA (Key Resources table) was identified using the CRISPOR design tool and purchased as a synthesized single guide RNA with modified ends from Synthego. One well of a 6-well plate at 70-80 % confluence was utilized for editing (approximately 10^6^ cells). Four hours prior to nucleofection, a full media change was performed with mTeSR Plus supplemented with Y-27632 [10 μM]. PSCs were then dissociated into single cells using TrypLE Express, and nucleofected using a Nucleofector 4D with the P3 Primary Cell kit following the manufacturer instructions (Lonza V4XP-3012) using 1 μg of Cas9 mRNA (Tri-Link, L-7206) and 2 μg of synthesized sgRNA. The nucleofected hPSCs were immediately neutralized with mTeSR Plus supplemented with Y-27632 (10 μM), and plated in a 10 cm dish pre-coated with Stem Cell Qualified BME Cultrex. The following day, the media was replaced with fresh mTeSR Plus media without Y-27632. Individual colonies that emerged were then manually transferred into 48-well plates for clonal expansion. Isogenic lines were screened by genomic DNA extraction, amplification of the NKX2-2 target locus by PCR (Key Resources table: Fwd NKX2.2, Rev NKX2.2) and characterization by Sanger sequencing. Two clones with a single base-pair deletion in the first exon of NKX2.2 were selected.

### METHOD DETAILS

#### hESC maintenance

MEL-1 INS^GFP/w^ cells were maintained as described by Hogrebe et al. Cells were seeded onto Cultrex-coated culture plates (Corning Cat# 0720083) in mTeSR Plus medium (StemCell Technologies cat. #100-0276) and passaged every 3 to 5 days. Aggregate passaging was performed by incubation in 0.5 mM EDTA for 10 minutes at 37 °C, resuspension of aggregates in mTeSR plus with 10 μM Y-27632, and seeded in new wells at a ratio of 1:20-40. After 24 hours, media was replaced with fresh mTeSR Plus. Media was then changed daily until the next passage. Maintenance cultures were discarded after 10 passages from thawing.

#### Directed differentiation

Cells were differentiated into endocrine progenitors and islet-like cells using the protocol of Hogrebe et al. with modifications. hESCs at 90 % confluence were dissociated to single cells with TrypLE Express, resuspended in mTeSR Plus with 10 μM Y-27632, and plated on Cultrex-coated plates at 3.125 x 10^5^ cells/cm^2^ for 24 hours. Cells were then washed with PBS and differentiated through six stages with stage-specific media. Medium was changed daily until day 21 and every other day thereafter.

#### Base media

All stage media were based on MCDB 131 supplemented with D-glucose, NaHCO3, Glutamax, BSA and penicillin-streptomycin. Stage 1: 9 mM D-glucose, 0.1 % BSA, 1 % Glutamax, 1 % penicillin-streptomycin, 14 mM NaHCO3. Stage 2: 4.5 mM D-glucose, 0.1 % BSA, 1 % Glutamax, 1 % penicillin-streptomycin, 14 mM NaHCO3. Stages 3 and 4: 2.4 mM D-glucose, 2 % BSA, 1 % Glutamax, 1 % penicillin-streptomycin, 21 mM NaHCO3. Stage 5: 20 mM D-glucose, 2 % BSA, 1 % Glutamax, 1 % penicillin-streptomycin, 21 mM NaHCO3. Stage 6: 2.4 mM D-glucose, 2 % BSA, 1 % Glutamax, 1 % penicillin-streptomycin.

#### Stage media-specific growth factor and small molecules

Stage 1 (4 days): 200 ng/mL Activin A, with 3 μM CHIR99021 on day 1 only. Stage 2 (2 days): 50 ng/mL KGF, 2.5 μM L-ascorbic acid. Stage 3 (2 days): 2 μM retinoic acid, 50 ng/mL KGF, 0.25 μM SANT1, 0.2 μM LDN, 0.2 μM TPPB, 2.5 μM L-ascorbic acid, 0.5 % ITS-X. Stage 4 (4 days): 0.1 μM retinoic acid, 50 ng/mL KGF, 0.25 μM SANT1, 0.2 μM LDN, 0.2 μM TPPB, 2.5 μM L-ascorbic acid, 0.5 % ITS-X. Stage 5 (7 days): 2 μM retinoic acid, 0.25 μM SANT1, 1 μM triiodothyronine, 1 μM γ-secretase inhibitor, 10 μM Alk5i II, 2.5 μM L-ascorbic acid, 0.5 % ITS-X, 10 mg/L heparin, with 1 μM latrunculin A on day 13 only. Stage 6 (14 days): 10 mg/L heparin, 1 % MEM non-essential amino acids, 0.1 % trace elements A, 0.1 % trace elements B, 1 μM ZnSO4 heptahydrate. Timepoints are named by stage and day, so S5d4 is the fourth day of stage 5 and S6d14 is the fourteenth day of stage 6.

#### Immunofluorescence

Cells were differentiated on glass coverslips, washed in PBS and fixed in 4 % paraformaldehyde for 30 minutes at room temperature. Cells were blocked and permeabilized in ICC solution (0.1 % Triton X-100 and 5 % normal donkey serum in PBS) for 45 minutes at room temperature. Primary antibodies (Supplemental Table 1) were diluted in ICC solution and applied overnight at 4 °C. Secondary antibodies were applied at 1:1000 in ICC solution for 2 hours at room temperature in the dark, followed by DAPI at 1:1000 for 12 minutes. Coverslips were mounted with ProLong Gold Antifade Mountant. Images were acquired on a Zeiss LSM 800 confocal laser scanning microscope and processed in Fiji (ImageJ v1.54p).

#### Flow cytometry

Cells were dissociated with TrypLE Express for 7 minutes at 37 °C, washed in PBS and fixed in 4 % paraformaldehyde for 12 minutes at RT. Cells were permeabilized in 1 X eBioscience permeabilization buffer (Invitrogen #00-8333-56) for 12 minutes at room temperature, then incubated overnight at 4 °C in primary antibodies (Supplemental table 1) diluted in CAS-Block with 0.4 % Triton X-100. Cells were washed in ICC solution. If secondary antibody staining was required, secondary antibody was then applied at 1:300 for 2 hours at 4 °C. Cells were washed, resuspended in 200 μL ICC solution and filtered. 50,000 cells per sample were acquired on a Cytek Aurora 5L spectral flow cytometer and analyzed in FlowJo v10.10.0. All samples were first gated to exclude debris, followed by doublet exclusion, and then analysis of marker expression (Supplemental Figure 5).

#### Bulk RNA sequencing

Cells were washed in PBS, dissociated with TrypLE Express for 8 minutes at 37 °C, pelleted at 300 x g for 3 minutes, washed in PBS, resuspended in RLT Plus buffer and frozen at −80 °C. RNA was isolated with the RNeasy Plus Mini Kit (Qiagen, cat no. 74134). Libraries were prepared by the CU Anschutz Genomics Shared Resource from at least 500 ng RNA using the Tecan Universal Plus mRNA-seq kit with NuQuant, and sequenced on an Illumina NovaSeq 6000 or NovaSeq X to approximately 40 million read pairs per sample. Reads were assessed with FastQC v0.12.1, adapters were trimmed with cutadapt v4.8, reads were aligned to hg38 with STAR v2.7.11b, and counts were generated with featureCounts from subread v2.0.6. Samples comprised three independent differentiations each of WT, NKX2.2KOc3 and NKX2.2KOc5 at S5d4 and at S6d14.

#### Single cell RNA sequencing

Cells were differentiated to S5d4, dissociated with TrypLE Express for 8 minutes at 37 °C, pelleted at 300 x g for 3 minutes and washed twice in PBS with 0.1 % BSA before delivery on ice to the CU Anschutz Genomics Shared Resource. Over 10,000 cells per sample were loaded onto a 10x Chromium controller for GEM formation and barcoding, and cDNA synthesis and library preparation were performed with the Chromium GEM-X Single Cell 3’ kit with v4 reagents. Library quality was assessed on a TapeStation (High Sensitivity D1000, Agilent) and libraries were sequenced on an Illumina NovaSeq X. 67,252 cells were captured and merged into the Seurat object. After QC filtering, doublet exclusion, and pancreatic subset selection 22,903 cells were retained.

#### Published NKX2.2 ChIP-seq data

Human islet NKX2.2 ChIP-seq peak files from two donors were obtained from ArrayExpress (E-MTAB-1919; Pasquali, et al.^42^). Peaks were called at p < 1 × 10⁻¹⁰ and are in hg19 coordinates. A reproducible peak set was defined as donor 1 peaks lying within 10 bp of a donor 2 peak (GenomicRanges subsetByOverlaps, maxgap = 10) giving 32,283 peaks. Peaks were annotated to the nearest gene with ChIPseeker v1.42.1^58^ using TxDb.Hsapiens.UCSC.hg19.knownGene and org.Hs.eg.db, with a promoter window of ±3 kb around the transcription start site. The nearest gene and the distance to the transcription start site were recorded for each peak.

Chromatin state was assigned by overlap with the human islet regulome (Supplementary Data Set 1 of Miguel-Escalada, et al.^43^) which classifies open chromatin regions by H3K27ac and Mediator signal. A peak was assigned to a regulome element if the two lay within 1 kb of each other. This tolerance was checked against the known MAFA enhancer as a positive control. Each peak received a single state in priority order: active promoter, active enhancer, inactive enhancer, CTCF, inactive open chromatin. Peaks overlapping no element were recorded as unannotated. The term inactive enhancer is used throughout in place of poised enhancer, because the classification is an H3K27ac call and does not include the bivalent marks that poised would imply.

### QUANTIFICATION AND STATISTICAL ANALYSIS

#### Bulk RNA sequencing differential expression

Counts were analyzed with DESeq2 v1.46.0^59^ in R. Genes with fewer than 10 reads summed across all 18 samples were removed, and a single DESeqDataSet spanning both timepoints was fit with a design of batch + genotype_day. That fit supplied the gene universe, the size factors and the variance stabilized values used for plotting. Each timepoint was then modeled separately by subsetting that object and refitting with a design of batch + genotype, so dispersions are estimated within the timepoint while size factors are retained from the shared fit. Batch is the sequencing run, and each batch included at least one sample from each cell line. Genotype contrasts the combined NKX2.2KO clones against WT, applied as a numeric contrast built from the column means of the model matrix for each genotype, so the two knockout clones are pooled and the batch terms cancel in this balanced design. Log fold change shrinkage was applied with ashr^60^, and genes with a Benjamini-Hochberg adjusted p below 0.05 were called differentially expressed, with no fold change threshold.

#### Bulk RNA sequencing figure panels

For the heatmaps, counts were variance stabilized with blind = FALSE and the sequencing run effect was removed with ComBat^61^ within each timepoint, with clone included in the ComBat model matrix so that genotype differences are preserved. Plotted genes were those called differentially expressed at that timepoint with an absolute log2 fold change above 1 and a mean normalized count of at least 5. Values were subset to that timepoint before scaling, rows were z-scored across the nine samples and clustered by complete linkage, and columns are shown in genotype order without clustering. Because the plotted genes were selected as differentially expressed against WT, the panels display the expression pattern of that set and are not evidence about the relationship between the two clones. A column-clustered version of each heatmap was generated as a quality control check and is not shown.

#### Selected-gene fold change panels

For these panels each knockout clone is tested against WT individually. Bars are the log2 fold change fitted by that model, error bars are its standard error, and WT is the reference at zero. An asterisk applies only to the bar above which it sits, and p-values are Benjamini-Hochberg adjusted. These estimates are not shrunk, so that the plotted value reflects the data for that gene directly. As a check, every plotted bar was compared against the mean ComBat-corrected variance stabilized difference from the WT mean for the same gene and samples, and any disagreement greater than one log2 unit was flagged.

#### Single cell processing, clustering and annotation

Alignment to hg38 and initial processing were performed with Cell Ranger v8.0.0, with each sample processed individually. Count matrices were read into Seurat^62^ and a Seurat object was created per sample. Low-quality cells were removed by filtering on number of features, number of counts and percentage of mitochondrial reads with perCellQCFilters from scuttle v1.8.4. Doublets were identified and removed with DoubletFinder v2.0.3^63^. WT and NKX2.2KO objects were merged, and batch correction across the six samples was performed by mutual nearest neighbor matching^64^ retaining 50 dimensions. Clustering and UMAP embedding were computed on the corrected reduction, with both genotypes clustered together. In total, 22,903 cells were retained across three independent differentiations per genotype. Clusters were annotated using the WT cells only. Each cluster was named from the significantly enriched expression of known population markers among its WT cells, giving 13 populations (Table 1, Supplemental Figure 2A). NKX2.2KO cells carry the label of the cluster they fall in. Cells of the two genotypes in the same cluster therefore share a transcriptional neighborhood but are not assumed to be equivalent. The replicate unit for all downstream statistics is the independent differentiation.

#### Pseudobulk differential expression

For each annotated population, raw counts were summed across cells within each independent differentiation to give one profile per population per replicate. Pseudobulk profiles built from fewer than 10 cells were excluded, and populations with fewer than two remaining replicates per genotype were not tested. Differential expression between genotypes was tested per population with DESeq2^59^, with log fold change shrinkage by apeglm^65^ and Benjamini-Hochberg correction, and genes with adjusted p below 0.05 were called differentially expressed.

#### Cell type proportion analysis

Population proportions were calculated for each independent differentiation as the percentage of that replicate’s cells assigned to each population. Genotypes were compared for each population by two-tailed Welch’s t-test on the three replicates per genotype. P-values were adjusted across all 13 populations with the Benjamini-Hochberg method, and adjusted p < 0.05 was considered significant. Bars show the mean of the three replicates, and points show individual replicates.

#### Co-expression analyses

Cells were classified as positive for a gene if normalized expression was above 0.5 for ARX, PAX4, NEUROG3, NKX2.2 and NKX6-1. For each analysis the percentage of cells in each class was calculated per population per independent differentiation and compared between genotypes by two-tailed Welch’s t-test. For the analysis of NEUROG3 and NKX2.2 co-detection along the WT progenitor trunk, cells were instead scored positive if any transcript was detected. Cells included in the ARX and PAX4 analysis all passed DoubletFinder filtering, and double-positive cells were additionally checked against the doublet call and retained as singlets. Note that NKX2.2 transcript remains detectable in the knockout, since the edit is a single base-pair deletion rather than a deletion of the locus, so the NKX2.2 classification is applied to WT cells only.

#### Trajectory inference and pseudotime

Trajectories were inferred with a single joint Slingshot v2.14.0^66^ fit on the first 10 dimensions of the batch-corrected reduction, with both genotypes pooled and the annotated populations as the cluster input. The fit was rooted at the bipotent progenitor population and terminated at the Alpha, Beta, Delta and Epsilon populations. The tree topology was fixed to the known developmental structure, with three arms leaving EP_early: EP_early to Alpha_prog to Alpha, EP_early to EP_late to Beta_prog to Beta, and EP_early to Delta_prog to Delta, with Epsilon attached to Alpha_prog. An unconstrained fit with the same start and endpoints reproduced the α, β and ε lineages but routed the δ lineage through Alpha_prog. Because δ fate is specified on the *PAX4*-dependent branch rather than the *ARX*-dependent α branch^47,67^ and routing δ through Alpha_prog would assign α progenitors to the δ lineage, the δ arm was fixed to branch from EP_early. Expression of *NEUROG3*, *HHEX*, *POU2F2* and *SST* along the δ lineage, and the WT versus NKX2.2KO comparisons in the late pseudotime window, were unchanged under the unconstrained route (Supplemental Figure 6A). Curves were fitted with reassignment disabled, so cells carry weight only on lineages that pass through their own population. Pseudotime values were extracted with slingPseudotime. Both genotypes were placed on the same curves, so WT and NKX2.2KO cells are compared on a common axis. For expression along pseudotime, each lineage was divided into 10 equal-width bins; within each bin, mean normalized expression was computed per independent differentiation and the plotted value is the mean of the three-replicate means with standard error across replicates. Cell counts per bin per genotype are plotted beneath each panel. For the late-pseudotime *NEUROG3* comparison, cells were scored *NEUROG3*-positive if any transcript was detected. The fraction of positive cells with scaled pseudotime of 0.65 or greater was computed per independent differentiation for each lineage, and genotypes were compared by two-tailed Welch’s t-test.

#### Branch allocation

A cell was counted as on a lineage if its Slingshot curve weight for that lineage was 0.5 or greater. Cells in a lineage’s progenitor or terminal population that were on that lineage alone were labeled as committed to it. Cells in the bipotent progenitor and EP_early populations were labeled progenitors. Cells in EP_late, which lies on the β lineage before Beta_prog, were labeled EP_late and not counted as committed β cells. Cells in Alpha_prog, which is shared by the α and ε lineages, were labeled Shared α/ε. For the density plot, each cell was placed at its mean pseudotime across the lineages it was on. The share of trajectory cells in each class was calculated per independent differentiation (Figure 6C), and the density of trajectory cells along pseudotime was computed per genotype with the same class labels (Figure 6B). The share of committed cells in each lineage was compared between genotypes by two-tailed Welch’s t-test on the three replicates per genotype. This analysis is reported alongside the cluster proportions because it measures position along the lineage and allocation between arms, which the cluster proportions do not.

#### Binding mode analysis

Genes were assigned a binding mode from the chromatin states of all NKX2.2 peaks annotated to them: active (active enhancer or active promoter only), inactive (inactive enhancer only), mixed (both active and inactive classes present) or unannotated only. For each mode the distribution of pseudobulk log2 fold changes was compared against genes with no peak, and modes were compared against each other, by two-sided Wilcoxon rank sum tests with Benjamini-Hochberg correction, both pooled across populations and per population.

#### Enrichment analysis

For each population, the set of genes that were both NKX2.2-bound and differentially expressed was tested for Gene Ontology Biological Process enrichment with gprofiler2^68^ v0.2.3, using the genes detected in that population as background and g:SCS correction. Terms with adjusted p below 0.05 were reported, and populations in which a term was not significantly enriched are shown as blank in Figure 4H rather than as a non-significant point. Overrepresentation analysis of the bulk differentially expressed genes was performed with the same package, restricted to significant terms.

#### General statistics

For all hESC experiments, n refers to the number of independent hESC differentiation experiments. Bar graphs show the mean with individual replicate values overlaid. Error bars represent the SEM unless stated otherwise. Two-sample comparisons of proportions and percentages used two-tailed Welch’s t-tests. Hormone class percentages from immunofluorescence were scored across three slides per condition and compared by two-way ANOVA. Differential expression significance is reported from DESeq2 Wald tests with Benjamini-Hochberg correction. Distributional comparisons of log fold change by binding mode used Wilcoxon rank sum tests with Benjamini-Hochberg correction. Statistical analyses were performed in R (v4.2.3 and v4.4.2, as indicated) and GraphPad Prism (v10.1.1). Statistical parameters and tests are reported in the figures and figure legends.

